# Scleraxis-lineage cells are required for tendon homeostasis and their depletion induces an accelerated extracellular matrix aging phenotype

**DOI:** 10.1101/2022.01.20.477119

**Authors:** Antonion Korcari, Anne EC. Nichols, Mark R. Buckley, Alayna E. Loiselle

## Abstract

Aged tendons have disrupted homeostasis, increased injury risk, and impaired healing capacity. Understanding mechanisms of homeostatic disruption is crucial for developing therapeutics to retain tendon health through the lifespan. Here, we developed a novel model of accelerated tendon extracellular matrix (ECM) aging via depletion of *Scleraxis-lineage (Scx*^*Lin*^) cells in young mice (DTR). DTR recapitulates many aspects of tendon aging including comparable declines in cellularity, alterations in ECM structure, organization, and composition. Single cell RNA-sequencing demonstrated a conserved decline in tenocytes associated with ECM biosynthesis in aged and DTR tendons, identifying the requirement for *Scx*^*Lin*^ cells during homeostasis. However, the remaining cells in aged and DTR tendons demonstrate functional divergence. Aged tenocytes become pro-inflammatory and lose proteostasis. In contrast, DTR tenocytes demonstrate enhanced remodeling capacity. Collectively, this study defines DTR a novel model of accelerated tendon ECM aging and identifies novel biological intervention points to maintain tendon function through the lifespan.

## Introduction

Tendons are dense connective tissues that transmit muscle-generated forces to bone to enable skeletal movement and joint stability. Recent studies from our group and others have demonstrated the presence of multiple tendon resident cell populations, with *Scleraxis-lineage (Scx*^*Lin*^) cells being the predominant population during adult mouse homeostasis [1-4]. *Scx*, a basic helix-loop-helix transcription factor, is the most well-characterized tendon marker [5-7], and is required for normal tendon development [5, 6]. During adulthood, tendon homeostasis is maintained via ongoing turnover of extracellular matrix (ECM) proteins [8-14]. Homeostatic disruptions can lead to the development of tendon pathologies such as tendinopathy, which are characterized by significant pain and permanent decline in tissue performance [15]. While repetitive loading in sports or occupational contexts disrupt homeostasis, natural aging is also associated with impaired tendon homeostasis [16-23]. Tendinopathy prevalence significantly increases with age [24], making aging a key risk factor for loss of tendon health. Despite this increased risk, the factors responsible for inducing age-related tendinopathy are still not well defined.

We have previously shown that depletion of *Scx*^*Lin*^ cells results in a relatively rapid loss of ECM structural and organizational integrity of the flexor digitorum longus (FDL) tendon [1], suggesting that *Scx*^*Lin*^ cells are required to maintain tendon homeostasis. However, the specific mechanisms by which *Scx*^*Lin*^ cells maintain tendon homeostasis have not been identified. Moreover, while the initial ECM organizational changes that occur with *Scx*^*Lin*^ cell depletion mimic those that occur during natural aging [17, 19, 25-27], it is unknown whether sustained *Scx*^*Lin*^ cell depletion in young mice may induce an aged tendon phenotype.

In the present study, our primary objective was to identify the underlying changes in the cellular and molecular environment that result from *Scx*^*Lin*^ cell depletion in young adult tendons and determine how these structural and compositional shifts compare to those observed during natural aging. Here, we define the mechanisms that are conserved between *Scx*^*Lin*^ depletion and natural aging to establish the key drivers that underpin the initiation of tendon degeneration and identify novel intervention points to maintain tendon health through the lifespan. Conversely, by defining the divergent molecular programming shifts in the tendon cells that remain during natural aging and inducible depletion of *Scx*^*Lin*^ tenocytes in young animals, we have identified cell populations and processes that may dictate differences in tendon healing capacity.

## Materials and Methods

### Mice

All animal studies were approved by the University Committee for Animal Resources (UCAR). Scx-Cre mice were generously provided by Dr. Ronen Schweitzer. ROSA26-iDTR^F/F^ (#007900) mice were obtained from the Jackson Laboratory (Bar Harbor, ME, USA). Scx-Cre mice were crossed to ROSA26-iDiphteria Toxin Receptor^Lox-STOP-Lox^ (DTR^F/F^) mice to generate a model of *Scx*^*Lin*^ tendon cell depletion (Scx-Cre^+^; DTR^F/+^; referred to as DTR). Expression of DTR is inhibited prior to Cre-mediated recombination due to the presence of a STOP cassette flanked by loxP sites (Lox-STOP-Lox). After Cre-mediated recombination, the STOP cassette is deleted, resulting in expression of DTR, specifically in *Scx*^*Lin*^ cells. Thus, administration of diphtheria toxin (DT) in these mice results in apoptosis of *Scx*^*Lin*^ cells. Scx-Cre^-^; DTR^F/+^ (WT) littermates were used as controls. DT was administered to both DTR and WT mice at three months of age. Male and female mice were used for all studies. C57BL/6J mice (B6; #664, Jackson Laboratory) were used for natural aging studies as noted. All mouse work (injections, surgeries, harvests) were performed in the morning. Mice were kept in a 12 hr light/dark cycle.

### Paraffin histology and immunofluorescence

Sections were stained with DAPI to visualize nuclei and imaged using a VS120 Virtual Slide Microscope (Olympus, Waltham, MA). Using ImageJ [28], a region of interest (ROI) was drawn at the tendon midsubstance and the area of the ROI was calculated. Nuclei within the ROI were manually counted and total nuclei number was normalized to area. An n = 5-7 mice per group were used for quantification. Five-micron sagittal sections of 6M WT, 6M DTR, and 31M C57BL/6J hindpaws were utilized For immunofluorescence, sections were stained GBP2 (1:500, Cat#: 11854-1-AP). Sections were counterstained with DAPI to visualize nuclei and imaged using a VS120 Virtual Slide Microscope (Olympus, Waltham, MA). *GBP2*^*+*^ cells and total nuclei within the ROI were manually counted and the number of *GBP2*^*+*^ cells was normalized by the total nuclei number to quantify the % of *GBP2*^*+*^ cells. All quantification was conducted in a blinded manner. An n = 6-8 mice per group were used for quantification.

### Second harmonic generation two-photon confocal imaging

Five-micron paraffin sections of DTR, WT, and C57BL/6J hind paws were utilized for second harmonic generation (SHG) imaging. Sections were scanned with a Spectra-Physics MaiTai HP DeepSee Ti:Sapphire Laser, tuned to 1000 nm, under 25x magnification, 2.5X optical zoom, and step size of 0.25 mm. 3D projections of image stacks were generated using the 3D-Project macro in ImageJ and analyzed for collagen fibril uniformity using the built-in Directionality macro function. The Directionality (dispersion) macro utilizes Fourier transform analysis to derive spatial orientation of image stacks [29]. Sections were analyzed from n=3-5 mice per group per age.

### Quantification of biomechanical properties

DTR and WT and C57BL6/J FDL tendons were harvested from the hind paws. Specifically, each FDL tendon was carefully separated at the myotendinous junction under a dissecting microscope. The tarsal tunnel was then cut and the FDL tendon was slowly released from the tarsal tunnel, isolated until the bifurcation of the digits and then cut and released. Under the dissecting microscope, any additional connective tissues (e.g., muscle) were removed and the FDL tendon was prepared for uniaxial testing. Two pieces of sandpaper were placed on each end of the tendon and glued together using cyanoacrylate (Superglue, LOCTITE). All the steps above were performed with the tissue periodically submerged in PBS to avoid any potential tissue drying. Each gripped tendon was transferred into a semi-customized uniaxial microtester (eXpert 4000 MicroTester, ADMET, Inc., Norwood MA). The microtester, along with the sample, was transferred to an inverted microscope (Olympus BX51, Olympus) to visualize the tendon and quantify the gauge length, width, and thickness. The gauge length of each sample was set as the end-to-end distance between opposing sandpaper edges and was set the same for all samples tested. The cross-section of the tendon was assumed to be an ellipse, where the width of the tissue represents the major axis and the thickness of the tissue represents the minor axis. Based on the optically measured width and thickness of each tendon, the area of the elliptical cross-section was computed. A uniaxial displacement-controlled stretching of 1% strain per second until failure was applied. Load and grip-grip displacement data were recorded and converted to stress-strain data, and the failure mode was tracked for each mechanically tested sample. The load-displacement and stress-strain data were plotted and analyzed to determine structural (*stiffness)* and material (*modulus*) properties. Specifically, the slope of the linear region from the load displacement graph was determined to be the stiffness of the tested sample. The slope of the linear region from the stress-strain graph was taken to equal the elastic modulus parameter of each tested tendon. Note that this calculation assumes that stress and strain are uniform within each specimen. A sample size of n=7 animals per group was utilized for biomechanical testing and analysis.

### Sample Preparation for Mass Spectrometry

Homogenization of each tendon tissue was performed by adding 150 uL of 5% Sodium Dodecyl Sulphate (SDS), 100 mM Triethylammonium bicarbonate (TEAB). Samples were vortexed and then sonicated (QSonica) for 5 cycles, with a 1 minute resting period on ice after each cycle. Lysates were then centrifuged at 15,000 x g for 5 minutes to collect cellular debris, and the supernatant was collected. Next, the total protein concentration was determined by bicinchoninic acid assay (BCA; Thermo Scientific), after which samples were diluted to 1 mg/mL in 5% SDS, 50 mM TEAB. A mass of 25 µg of protein from each sample was reduced with dithiothreitol to 2 mM, followed by incubation at 55°C for 60 minutes. Iodoacetamide was added to 10 mM and incubated for 30 minutes in the dark at room temperature to alkylate the proteins. Phosphoric acid was added to 1.2%, followed by six volumes of 90% methanol, 100 mM TEAB. The resulting solution was added to S-Trap micros (Protifi) and centrifuged at 4,000 x g for 1 minute. S-Traps with the trapped protein were washed twice by centrifuging through 90% methanol, 100 mM TEAB. 1 ug of trypsin was brought up in 20 μL of 100 mM TEAB and added to the S-Trap, followed by 20 µL of TEAB to ensure the sample did not dry out. The cap to the S-Trap was loosely screwed on but not tightened to ensure the solution was not pushed out of the S-Trap during digestion. Samples were placed in a humidity chamber at 37°C overnight. The next morning, the S-Trap was centrifuged at 4,000 x g for 1 minute to collect the digested peptides. Sequential additions of 0.1% Trifluoroacetic acid (TFA) in acetonitrile and 0.1% TFA in 50% acetonitrile were added to the S-trap, centrifuged, and pooled. Samples were frozen and dried down in a Speed Vac (Labconco), then re-suspended in 0.1% trifluoroacetic acid prior to analysis. Three tendons per genotype per timepoint were used for proteomic analysis.

### Mass Spectrometry (MS)

Peptides were injected onto a homemade 30 cm C18 column with 1.8 um beads (Sepax), with an Easy nLC-1200 HPLC (Thermo Fisher), connected to a Fusion Lumos Tribrid mass spectrometer (Thermo Fisher). Solvent A was 0.1% formic acid in water, while solvent B was 0.1% formic acid in 80% acetonitrile. Ions were introduced to the MS using a Nanospray Flex source operating at 2 kV. The gradient began at 3% B and held for 2 minutes, increased to 10% B over 6 minutes, increased to 38% B over 95 minutes, then ramped up to 90% B in 5 minutes and was held for 3 minutes, before returning to starting conditions in 2 minutes and re-equilibrating for 7 minutes, for a total run time of 120 minutes. The Fusion Lumos was operated in data-dependent mode, with MS1 scans acquired in the Orbitrap, and MS2 scans acquired in the ion trap. The cycle time was set to 2 seconds. Monoisotopic Precursor Selection (MIPS) was set to Peptide. The full scan was done over a range of 375-1400 m/z, with a resolution of 120,000 at m/z of 200, an AGC target of 4e5, and a maximum injection time of 50 ms. Peptides with a charge state between 2-5 were picked for fragmentation. Precursor ions were fragmented by collision-induced dissociation (CID) using a collision energy of 30% with an isolation width of 1.1 m/z. The Ion Trap Scan Rate was set to Rapid, with a maximum injection time of 35 ms, an AGC target of 1e4. Dynamic exclusion was set to 45 seconds.

### MS data filtering

Raw data was searched using the SEQUEST search engine within the Proteome Discoverer software platform, version 2.4 (Thermo Fisher), using the SwissProt mus musculus database. Trypsin was selected as the enzyme allowing up to 2 missed cleavages, with an MS1 mass tolerance of 10 ppm, and an MS2 mass tolerance of 0.6 Da.

Carbamidomethyl was set as a fixed modification, while oxidation of methionine was set as a variable modification. The Minora node was used to determine relative protein abundance between samples using the default settings. Percolator was used as the FDR calculator, filtering out peptides which had a q-value greater than 0.01.

### Filtered MS data analysis

To identify proteins that were significantly different between two groups, the log2 fold change (FC) and the - log10 of the p-value were plotted in a volcano plot and we utilized the most stringent cut-off parameters of log2FC> 1 and -log10(p-value) >1.3. Significantly decreased proteins among two groups were inserted in PANTHER classification system [30] (http://pantherdb.org/) to classify protein type. To understand what types of biological processes (BPs), molecular functions (MFs), and cellular components (CCs) were shifted on each condition, we inserted all significantly decreased proteins in the gene ontology classification tool DAVID [31] (https://david.ncifcrf.gov/). Protein network analysis was performed using the Search Tool for Retrieval of Interacting Genes/Proteins (STRING), v11.0 [32], and ECM-related proteins were further classified using MatrisomeDB [33] (https://web.mit.edu/hyneslab/matrisome/).

### Single cell isolation

A total of 16 FDL tendons per group per age were pooled together into low glucose Dulmecco’s Modified Eagle Medium (DMEM) and subsequently digested in Collagenase Type I (5mg/ml) (Worthington Biochemical, Lakewood, NJ, LS004196) and Collagenase Type IV (1mg/ml) (Worthington Biochemical, Lakewood, NJ, LS004188). Young WT tendons were completely digested after two hours, while young DTR and aged C57BL/6J WT tendons after ninety minutes. After digestion, the single-cell suspension was filtered for any potential debris via a 70 μm and a 50 μm cell strainer and resuspended in Dulbecco’s Phosphate Buffer Solution (dPBS). Next, the single cell suspension was centrifuged for 10min at 500xg while being at 4°C. After removal of supernatant, the single cell suspension was resuspended in 0.5% Bovine Serum Albumin (BSA).

### Single cell RNA-sequencing and analysis

Single cell RNA-sequencing (scRNA-seq) was performed in collaboration with the UR Genomics Research Center (GRC). Libraries were prepared using a Chromium Single Cell 3 Reagent Kit (version 3, 10X Genomics, Pleasanton, CA) following the directions of the manufacturer. Cells were loaded into the kit and processed using a chromium controller (10X Genomics). Following library preparation and quality control (QC), libraries were sequenced using the S2 NovaSeq flow cell system (Illumina, San Diego, CA), generating an average of 65,000 reads per cell for all groups. Raw data quality control and downstream analyses were performed with R-4.1.2 and the Seurat package (4.0.6) [34]. Cells with < 1000 or > 5000 genes, as well as cells with > 10% mitochondrial genes were removed from the dataset. After filtering out low quality cells, there were 8,359 cells for the young (6M old) WT group, 6,554 cells for the young (6M old) DTR group, and 4,798 cells for the old (21M old) C57BL/6J WT group. All three different datasets were integrated to determine alterations of different cell subpopulations between groups using the “FindIntegrationAnchors” function (dims = 1:20). The function “IntegrateData” was applied in the anchor set with the default additional arguments. Next, data was scaled using the “ScaleData” with default parameters and principal component analysis (PCA) was performed on the integrated dataset to compute 20 principal components (PCs). Uniform Manifold Approximation and Projection (UMAP) dimensionality reduction was applied and the Shared Nearest Neighbour (SSN) graph was constructed by utilizing dimensions 1:20 as input feature. To identify cell clusters, the “FindClusters” function was utilized on the integrated dataset at a resolution of 0.1, resulting in seven distinct cell clusters. To identify genetic markers of each single cluster and annotate the different clusters to known cell types, the top differentially expressed genes of each cluster were identified in the heatmap ***(Fig. S4)***. Next, the “FindAllMarkers” function was used to identify upregulated genes for each cluster. The average gene expression level was calculated across each cluster. The minimum percentage of cells in which the gene is detected in each cluster was set to 50%. The average log2 change threshold was set to at least 0.25. Marker gene lists were then compared to a set of literature-defined gene markers via direct literature search and by utilizing *CellMarker* software (http://bio-bigdata.hrbmu.edu.cn/CellMarker/search.jsp) [35]. Significance was determined using Wilcoxon rank sum test with p values adjusted based on Bonferroni correction applying all features in the data set (p_val_adj<0.05). The tenocytes cluster from the integrated data was subset out from the rest using the “Subset” and were independently re-clustered following the same steps described above.

### Cell-cell communication analysis

Cellular autocrine and paracrine communication was performed using the R *CellChat* package version 1.4.0 (https://github.com/sqjin/CellChat). To assess tenocyte-tenocyte communication, tenocyte clusters 1-3 from the integrated data were subsetted and a CellChat object of tenocytes 1-3 was created for each condition using the “createCellChat” function. After proper annotation, the “CellChatDB.mouse” database was utilized for our study To identify overexpressed genes, the “identifyOverExpressedGenes” function was utilized. To identify significant cellular interactions, the “identifyOverExpressedInteractions” function was utilized. Finally, to infer communication probabilities, the “computeCommunProb” function was utilized. Next, to study the effect of depletion and natural aging on tenocyte-tenocyte communication, a merged CellChat object was created for the 6M WT vs 6M DTR, and 6M WT vs 21M B6 groups, respectively, using the “mergeCellChat” function. To assess tenocyte-macrophage communication, tenocytes and macrophage clusters from the integrated data were subsetted. The same steps mentioned above for the tenocyte-tenocyte communication analysis were also followed for tenocyte-macrophage communication. To identify the differential number of interactions between two conditions, the “netVisual_diffInteraction” function was utilized. To identify specific signaling pathways that were conserved or expressed only in one condition, the “rankNet” function was utilizes. Next, the “netAnalysis_signalingRole_heatmap” function were applied to identify the “outgoing” and “incoming” patterns of signaling pathways for each different cell population and how that compares between 2 conditions. Finally, specific ligand-receptor interaction were identified for specific signaling pathways of interest utilizing the “netVisual_bubble” function.

### Statistical analysis

Sample sizes for cell density, dispersion, and biomechanical properties were determined based on post-hoc power calculations of previously published work [3, 36]. Quantitative data was analyzed via GraphPad Prism and is presented as mean ± standard deviation (SD). Either a student’s t-test or two-way analysis of variance (ANOVA) with Tukey’s test was used as appropriate. Mice were randomly selected for specific experimental outcome metrics prior to the start of an experiment or surgery and quantitative data (ex. DAPI quantification, SHG dispersion levels, biomechanical properties) were analyzed in a blinded manner. For all experiments, an n=1 represents one mouse. p values ≤ 0.05 were considered significant. * indicates p<0.05, ** indicates p<0.01, *** indicates p<0.001, **** indicates p<0.0001.

## Results

### *Scx*^*Lin*^ cell depletion in young adults disrupts tendon homeostasis and mimics the cell density and ECM structural alterations of tendon aging

To determine the long-term depletion efficiency of this DTR model, we quantified cell density in DTR mice at 3-, 6-, and 9-months (3, 6, 9M) post-depletion, and compared to age-matched DT-treated WT littermates (***Figure 1A***). At 3M and 6M post-depletion, there was a respective 57.44% (p<0.0001) and 56.21% (p<0.0001) reduction in total tendon cell density in DTR tendons relative to the WT littermates (***Figure 1B, C***). By 9M post-depletion (12M of age), there was an age-related decrease in tendon cell density in WT such that no significant differences in cell density were observed between DTR and WT (p>0.05) (***Figure 1B, C)***.

**Figure 1.**
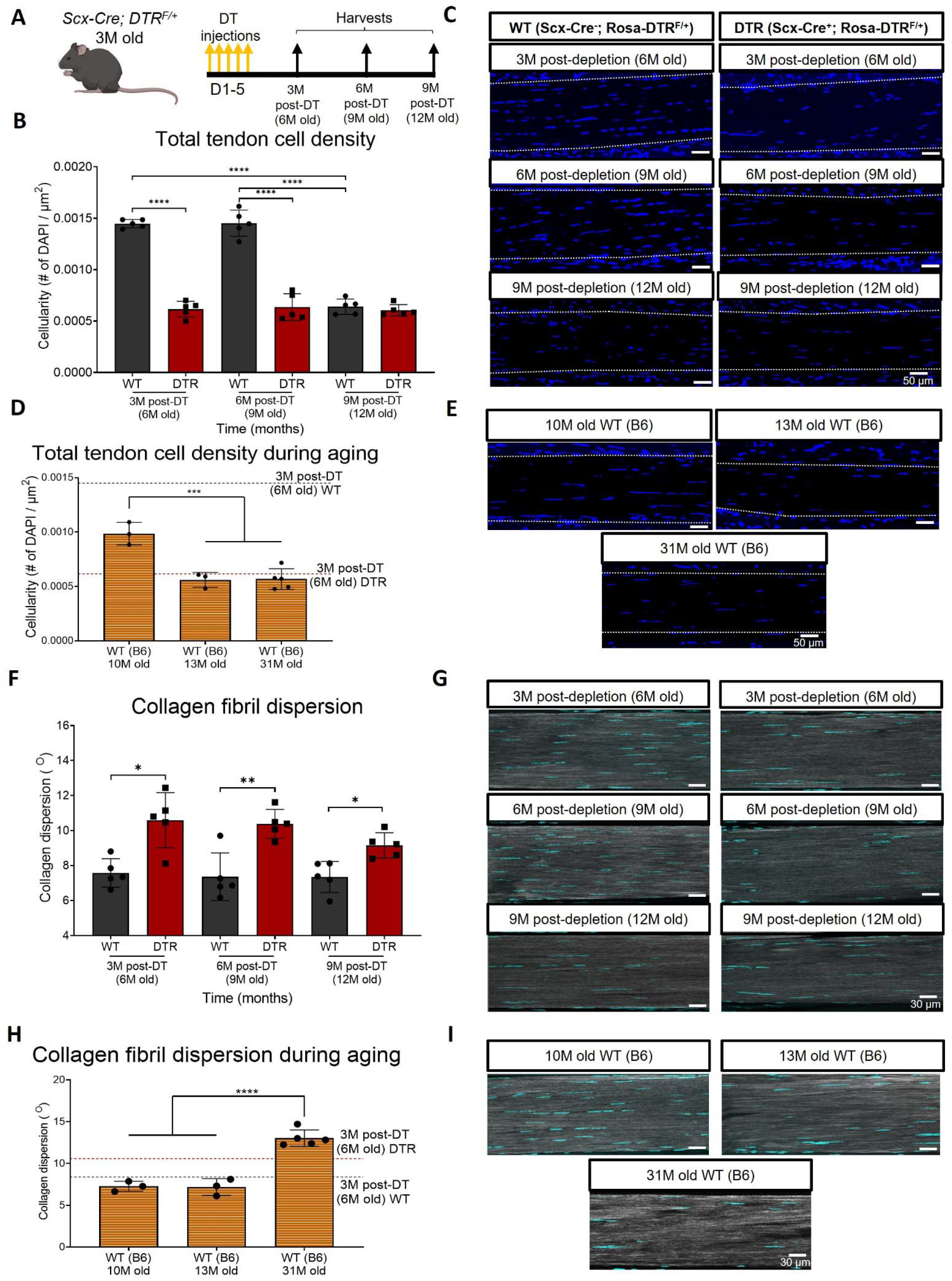
Depletion of *Scx*^*Lin*^ cell during long-term homeostasis significantly disrupts tendon structure and mechanical properties. **(A)** 3M old Scx-Cre; DTR^F/+^ mice received five hindpaw injections of DT and were harvested at 3, 6, and 9M post-depletion. **(B)** Quantification of total tendon cell (DAPI) density from injected WT and DTR hindpaws at 3, 6, and 9 months post-depletion (6, 9, and 12 months old of age, respectively) within the tendon. **(C)** Representative sections from **(B). (D)** Quantification of total tendon cell (DAPI) density from C57BL/6J hind paws at 10, 13, and 31 months old within the tendon. **(E)** Representative sections from **(D). (F)** Quantification of collagen fibril dispersion in WT and DTR samples at 3, 6, and 9 months post-depletion (6, 9, and 12 months old, respectively) and **(G)** representative collagen fibril morphology captured via SHG. **(H)** Quantification of collagen fibril dispersion from C57BL/6J hind paws at 10, 13, and 31 months old and **(I)** representative collagen fibril morphology via SHG. N=3-5 per genotype.

Based on this decline in cellularity from 9-12M in WT, we further tracked changes in tendon cell density from 10-31 months of age in C57BL/6J mice ***(Figure 1D)***. Consistent with the progressive decline in cellularity in WT tendons, cellularity was further decreased at 10M in C57Bl/6J tendons, with a 31.62% (p<0.0001) decline in cell density compared to 6M old WT, and a 32.06% decrease (p<0.0001) compared to 9M old WT tendons ***(Figure 1D, E)***. At 13 and 31M, C57Bl/6J FDL tendons showed a 43.19% (p<0.001) and a 44.42% (p<0.001) decrease in total cell density compared to 10M old C57Bl/6J FDL tendons, respectively ***(Figure 1D, E)***. Intriguingly, 6M old DTR tendons (3M post-depletion) exhibited a cell density almost identical to 13M and 31M old C57Bl/6J tendons (p>0.05) ***(Figure 1D, E)***, suggesting that with *Scx*^*Lin*^ depletion, young tendons exhibit the same cell density as old (12M) and geriatric (31M) tendons ***(Figure 1D, E)***.

We then investigated the impact of sustained *Scx*^*Lin*^ cell loss on long-term maintenance of collagen ECM organization. Quantification of collagen dispersion via SHG imaging demonstrated significant increases in dispersion (loss of organization) at 3M (39.82%, p<0.05), 6M (41.04%, p<0.01), and 9M (24.53%, p<0.05) post-DT, relative to WT littermates (***Figure 1F, G***), indicative of impairments in tissue structure and organization.

We then determined how the structure and organization of the tendon shifts with natural aging, and whether there may be conserved structural changes with both *Scx*^*Lin*^ depletion and aging. Geriatric 31M old C57Bl/6J tendons exhibited a significant increase in collagen fibril dispersion compared to 10M (79.3%; p<0.0001) and 13M (81.47%, p<0.0001) C57Bl/6J tendons ***(Figure 1H, I)***, suggesting aging-induced impairments in tendon structure and organization. No differences in dispersion were observed in 3M post-DT (6M old) WT tendons compared to 10M C57BL/6J (p>0.05), and 13M C57BL/6J (p>0.05) ***(Figure 1H, I)***. In contrast, 3M post-DT (6M old) DTR tendons showed a 45.8% (p<0.01) and a 47.63% (p<0.01) increase in collagen dispersion compared to 10M and 13M C57BL/6J tendons respectively ***(Figure 1H, I)***, suggesting that young DTR tendons recapitulate several aspects of age-related deficits in tissue organization. However, geriatric 31M old C57BL/6J tendons still had a significantly higher collagen fibril dispersion (+23.02%, p<0.05) compared to 3M post-DT (6M old) DTR tendons ***(Figure 1H, I)***, potentially suggesting further degeneration during advanced aging.

### *Scx*^*Lin*^ cells maintain tendon homeostasis by regulating the synthesis of high turnover rate ECM proteins

We next sought to identify the specific biological mechanisms that accompany, and potentially underpin these structural changes by characterizing the proteome of DTR and WT tendons at 3M post-depletion (6M old), as well as the proteome of old vs young WT tendons (12M and 6M old). At 3M post-depletion, 30 proteins were significantly different between the DTR and WT groups (***Figure 2A***). ECM proteins were the most decreased category, accounting for 26.7% of the total proteins. (***Figure 2B)***. In addition, cytoskeletal proteins, transporters, and scaffold/adaptor proteins were decreased, though these decreases in inter- and intra-cellular proteins are likely due to a ∼60% reduction in cellularity of the DTR samples. Functional enrichment analysis of all the downregulated proteins demonstrated that ECM-related molecular functions and cellular components were significantly impaired with depletion (***Figure 2C)***. Eleven ECM proteins were decreased in DTR tendons (***Figure 2D)***, and they were classified as proteoglycans (PG) (36.4%), glycoproteins (GP) (36.4%), ECM regulators (18.2%), and ECM-affiliated proteins (9.1%) (***Figure 2E)***. Strikingly, 72.8% of these ECM proteins were high turnover rate glycoproteins and proteoglycans including CHAD, COCH, and KERA [8, 10, 11, 37]. Similar findings were also identified at the 9M post-depletion timepoint (12M old) ***(Figure S1A-F)***.

**Figure 2.**
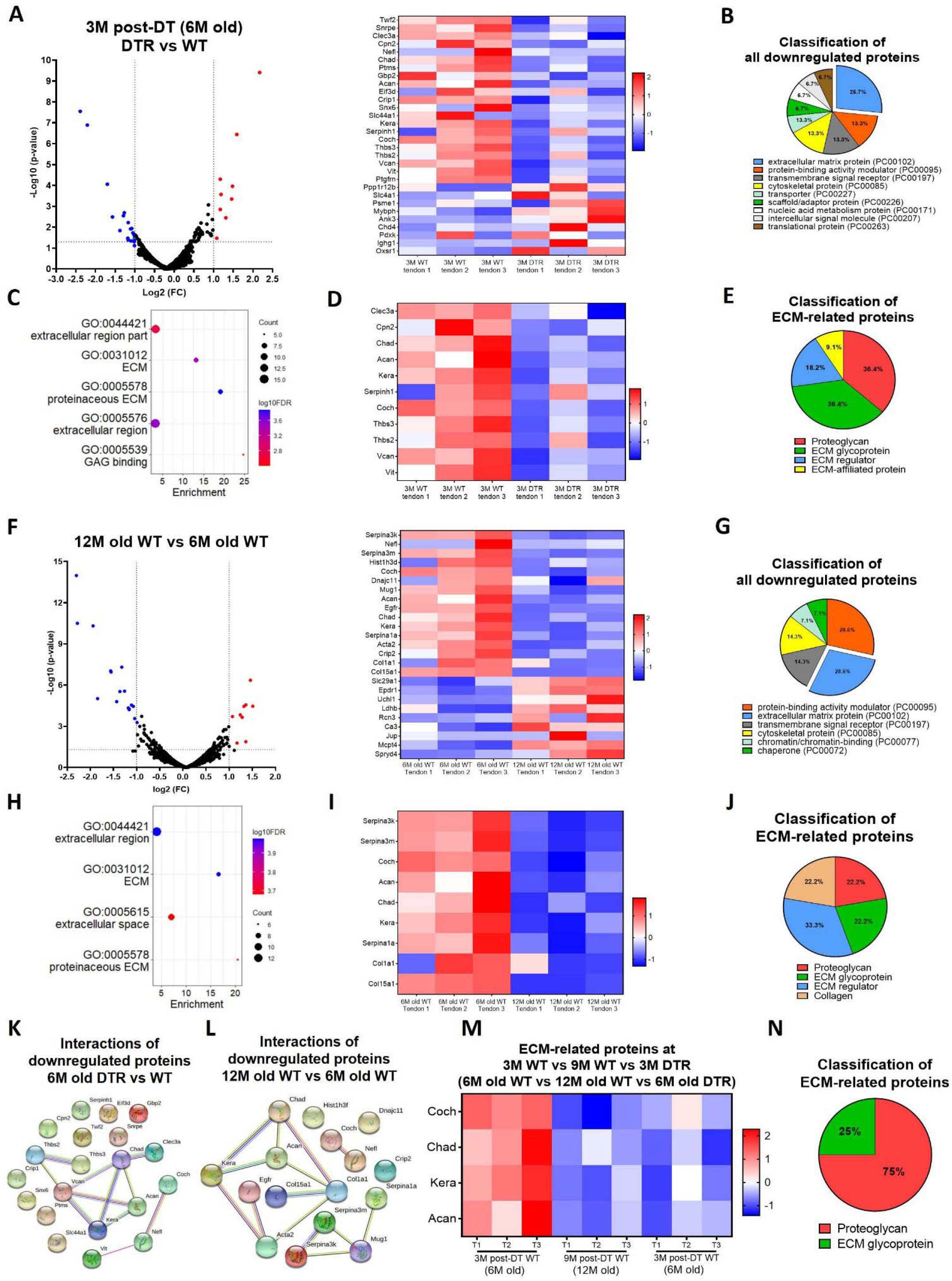
*Scx*^*Lin*^ cells maintain FDL tendon homeostasis by regulating the synthesis of high turnover rate ECM proteins. **(A)** Volcano plot and heatmap visualizing the significantly different protein abundances between DTR and WT groups at 3M post-depletion. **(B)** Classification of all downregulated proteins between the DTR and WT FDL tendons at 3M post-depletion. **(C)** Functional enrichment analysis of cellular components and molecular functions of all downregulated proteins between the DTR and WT FDL tendons at 3M post-depletion. **(D)** Heatmap of all differentially abundant ECM-related proteins between the DTR and WT FDL tendons at 3M post-depletion. **(E)** Classification of all ECM-related downregulated proteins between the DTR and WT FDL tendons at 3M post-depletion. **(F)** Volcano plot and heatmap visualizing the significantly different protein abundances between 12M and 6M WT FDL tendons. **(G)** Classification of all downregulated proteins between 12M and 6M WT FDL tendons. **(H)** Functional enrichment analysis of all downregulated proteins between 12M and 6M WT FDL tendons. **(I)** Heatmap of all differentially abundant ECM-related proteins between 12M and 6M WT FDL tendons. **(J)** Classification of all ECM-related downregulated proteins between 12M and 6M WT FDL tendons. **(K)** Protein-protein interaction of all the downregulated proteins between DTR and WT groups at 3M post-depletion. **(L)** Protein-protein interaction of all the downregulated proteins between 12M and 6M WT FDL tendons. **(M)** Heatmap with 4 ECM-related proteins that were decreased in similar rates with both natural aging and *Scx*^*Lin*^ cell depletion. **(N)** Classification of the ECM-related proteins from **(M)**.

With natural aging, 25 proteins were significantly different between the aged (12M) and young (6M) WT tendons ***(Figure 2F)***. Classification of all downregulated proteins determined that 28.6% were ECM proteins ***(Figure 2G)*** and functional enrichment analysis also showed that ECM-related molecular functions and cellular components were significantly impaired ***(Figure 2H)***. A total of 9 decreased ECM proteins were identified ***(Figure 2I)***, and classified as proteoglycans (22.2%), glycoproteins (22.2%), ECM regulators (33.3%), and collagens (22.2%) ***(Figure 2J)***. The majority of the above ECM molecules have a high turnover rate, similar to the findings for the 3M post-depletion timepoints ***(Figure 2D, E)***. Finally, the proteomes of aged (12M old) WT tendons vs young (6M old) DTR were nearly identical ***(Figure S2A, B)***. Collectively, these data further support the hypothesis that *Scx*^*Lin*^ cell depletion in young tendons mimic impairments in ECM-related biological mechanisms that occur during natural aging.

String DB found that in DTR tendons there are two separate groups of downregulated proteins ***(Figure 2K)***, suggesting two separate mechanisms of ECM-degradation with *Scx*^*Lin*^ depletion. With aging, three separate groups of proteins were downregulated ***(Figure 2L)***. Interestingly, proteins CHAD, ACAN, and KERA, as well as COCH and NEFL were identified also in the *Scx*^*Lin*^ depleted vs WT ***(Figure 2K)***, suggesting that these two mechanisms are shared with both natural aging and depletion. Based on these analyses, we identified four ECM molecules (COCH, CHAD, KERA, and ACAN) that were consistently downregulated with both natural aging and *Scx*^*Lin*^ cell depletion ***(Figure 2M)***. Classification showed that 75% are proteoglycans and 25% are glycoproteins ***(Figure 2N)***. Considering the substantial changes in ECM organization and composition, we also assessed the impact of *Scx*^*Lin*^ cell depletion and natural aging on tendon mechanical integrity (***Fig. S3A-D***) and while there were no major changes in CSA ***(Fig. S3B)*** and stiffness ***(Fig. S3B)***, there was a significant decrease of tendon elastic modulus with both depletion and natural aging ***(Fig. S3D)***.

### Single cell RNA-seq demonstrates similar loss of ECM biosynthetic tenocytes as well as differential retention of ECM organizational and age-impaired tenocytes in DTR vs aged tendons

Given that *Scx*^*Lin*^ cell depletion in young animals induced similar declines in cell density and ECM structure-function, as well as overlapping alterations in ECM composition compared to aged WT tendons, we next asked whether the composition of the cellular environment was conserved in the context of depletion vs. natural aging. Therefore, we performed scRNA-seq in 6M old WT and DTR tendons (3M post-DT) and 21M old C57BL/6J (B6) tendons. 21M old B6 tendons were used as the ‘aged’ group as we have previously established that by this age there are age-related impairments in tissue homeostasis and tendon healing response [38].

In the integrated dataset, seven cell types with distinct transcriptomic signatures were identified, including tendon fibroblasts (tenocytes), epitenon cells, endothelial, nerve, and muscle cells, macrophages, and T cells ***(Figure 3A, B; Fig. S4)***. Annotation of the epitenon cell cluster is based on additional work from our laboratory that will be published in a separate report ***(Fig. S4)***. Considering that *Scx*^*Lin*^ cells are tissue-resident fibroblasts (tenocytes), and these are the cells that are subject to depletion, our primary analysis focused on the impact of depletion and aging on tenocytes. We subsetted and re-clustered the tenocyte population ***(Figure 3C, D)*** from the integrated dataset ***(Figure 3A)***. Consistent with cellularity quantification ***(Figure 1)***, 6M DTR and 21M B6 tendons had a decrease in the total number of tenocytes captured compared to 6M WT ***(Figure 3D, E)***. To decipher the biological functions of the tenocytes and assess any potential intrinsic programmatic shifts resulting from depletion or natural aging, we investigated their unique transcriptomic profiles by identifying the differentially expressed genes (DEG) between the 6M WT, 6M DTR, and 21M B6 groups ***(Figure 3F)***, and performing functional enrichment analysis (FEA) based on their top 100 DEG ***(Figure 3 G-I)***.

**Figure 3.**
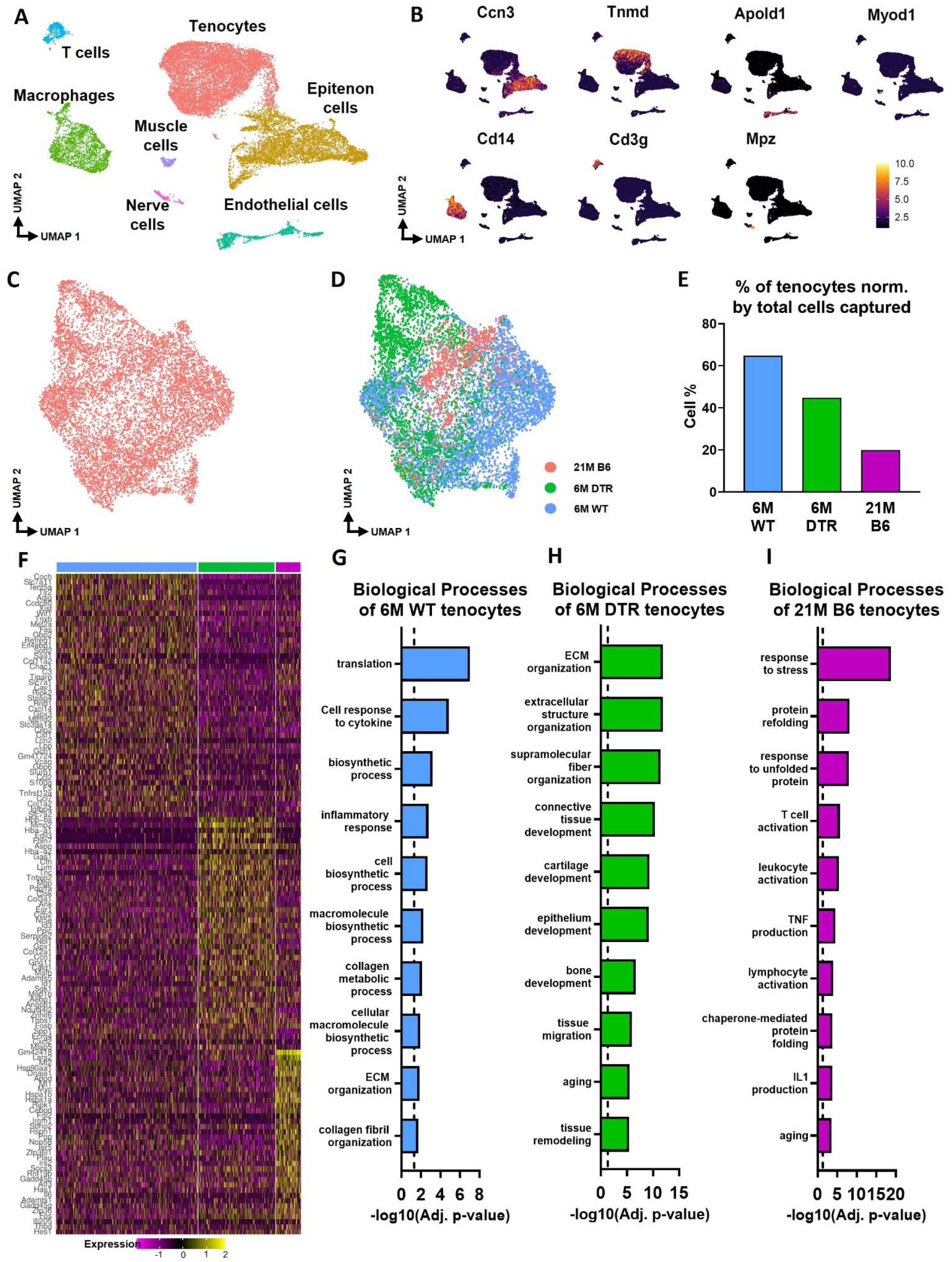
scRNA-seq demonstrates broad cellular heterogeneity and intrinsic programmatic skewing of tenocytes with depletion and natural aging. **(A)** UMAP dimensionality reduction revealed 7 broad and distinct cell populations on clustering based on unbiased differential gene expression of the integrated dataset. **(B)** Annotation marker for each of the 7 cell populations identified in the FDL tendons. **(C)** Re-clustering of the tenocytes cell population from **(A). (D)** UMAP plot of tenocytes in the integrated data colored based on respective group (Red: 21M old B6, green: DTR 6M old, blue: WT 6M old). **(E)** Quantification of the percentage of tenocytes normalized by the total cells captured per condition. **(F)** Heatmap with the top 40 significantly expressed genes in tenocytes per condition. **(G)** Significantly upregulated biological processes of tenocytes in the 6M WT **(G)**, 6M DTR **(H)**, and 21M B6 **(I)**. Dotted lines in **G, H**, and **I** indicate statistical significance of adjusted p-value<0.05 (-log10(adj. p-value = 1.3).

Tenocytes in the 6M WT group express genes related primarily to ECM biosynthetic process and secondary to ECM organization such as *Col1a1, Col1a2, Tnxb, Coch, Col11a2, and Fbln1*. Based on FEA, 6M WT tenocytes are broadly characterized as ECM biosynthetic (“translation”, “macromolecule biosynthetic process”, “collagen metabolic process”) and ECM organizational (“ECM organization”, “homeostatic process”, “collagen fibril organization”) ***(Figures 3F, G)***. Consistent with our proteomics findings, broadly, both DTR and aged tenocytes exhibited a decrease in processes related to ECM biosynthesis such as “translation”, “biosynthetic process”, “cell and macromolecule biosynthetic process” ***(Figure 3F-I)***. To better understand tenocyte heterogeneity as well as the similar shifts of tenocyte biological processes in DTR and natural aging, we subsetted and re-clustered the tenocyte cluster, which resulted in three distinct subpopulations ***(Figure 4A)***, and then examined the distribution of these subsets among the different groups (6M WT, 6M DTR, 21M B6) ***(Figure 4B-E)***. With both depletion and aging, we found that tenocytes 1 were decreased to almost identical levels.

**Figure 4.**
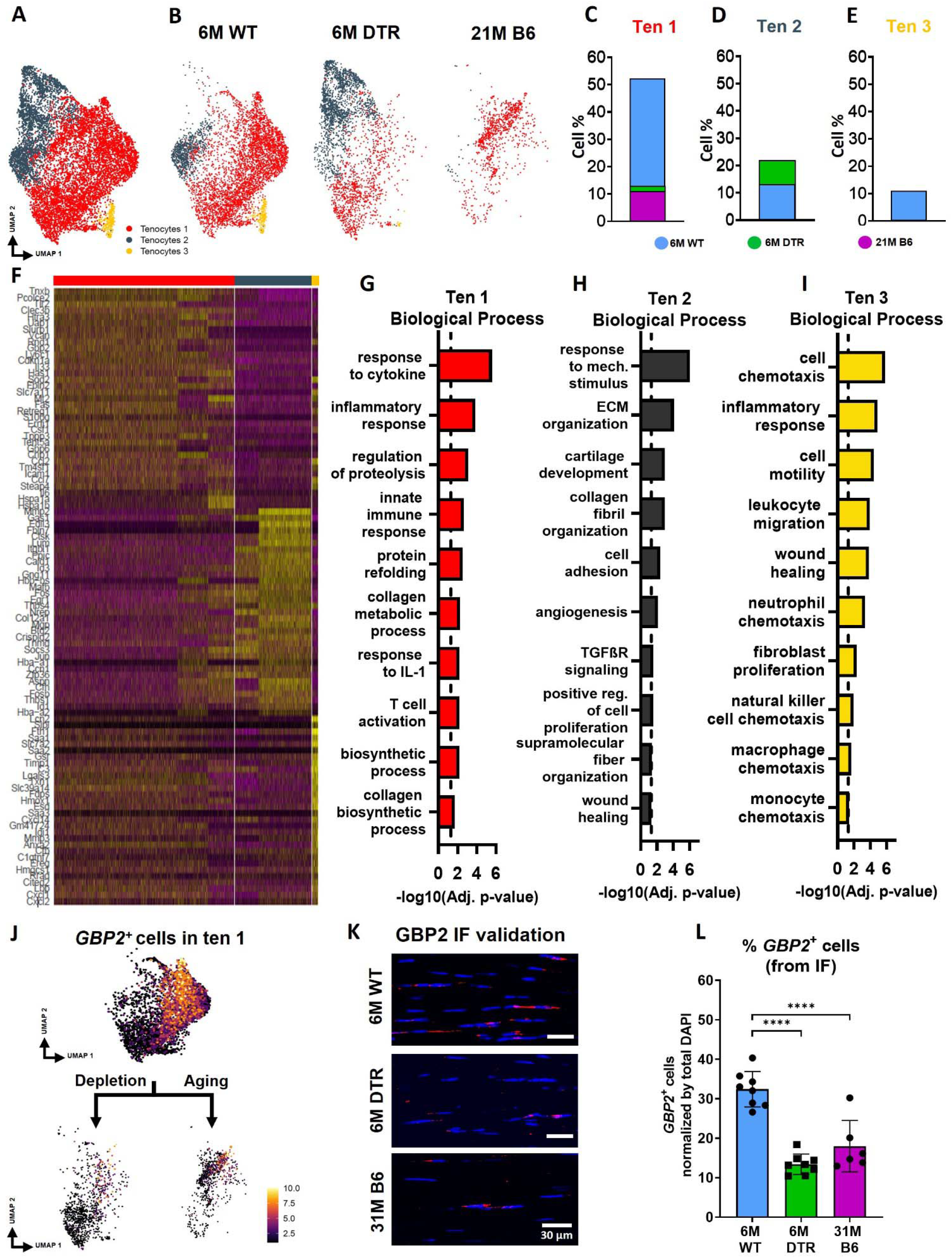
DTR and aged tendons lose tenocytes associated with ECM biosynthesis and immune surveillance, while DTR tendons retain tenocytes associated with ECM organization and remodeling. **(A)** UMAP plot of all three tenocytes subpopulations. **(B)** UMAP plot showing the shifts of tenocytes 1-3 with depletion and natural aging. **(C-E)** Quantification of % in tenocytes 1 **(C)**, tenocytes 2 **(D)**, and tenocytes 3 **(E)**, in the 6M WT, DTR, and 21M B6 groups **(F)** Heatmap with the top 40 significantly expressed genes per tenocyte subcluster. **(G-I)** Significantly upregulated biological processes of tenocytes 1 **(G)**, tenocytes 2 **(H)**, and tenocytes 3 **(I)**, respectively. **(J)** UMAP feature plot of *GBP2*^*+*^ tenocytes and their shift with depletion and natural aging, respectively. **(K)** Representative IF of protein validation on the decrease in *GBP2*^*+*^ tenocytes 1 cells with *Scx*^*Lin*^ cell depletion and natural aging. **(L)** Quantification of IF for *GBP2*^*+*^ cell density in 6M WT, 6M DTR, and 31M B6 groups.

Specifically, tenocytes 1 subpopulation exhibited a 75.18% decrease with depletion and a 78.9% decrease with aging ***(Figure 4B-E)***. FEA analysis demonstrated that tenocytes 1 express biological processes related to ECM biosynthesis (“biosynthetic process”, “collagen biosynthetic” and “collagen metabolic process”) and immune response (“inflammatory” and “immune response”, “response to IL1”, “T cell activation”) ***(Figure 4F, G)***. Finally, we identified *Gbp2* as a specific marker of tenocytes 1 ***(Figure 4J***) and found that with both depletion and natural aging there was a decrease in *Gbp2*^*+*^ cells compared to the 6M WT ***(Figure 4J)***. Staining for GBP2 protein validated our scRNAseq results and showed a 58.66% *(p<0*.*0001)* and a 44.54% *(p<0*.*0001)* decrease in *GBP2*^*+*^ cells with depletion and natural aging relative to the 6M WT group ***(Figure 4K, L)***. Taken together, our scRNAseq results demonstrate that the consistent loss of ECM biosynthetic tenocytes 1 in young DTR and aged groups is the main driver for impaired tendon homeostasis.

Besides the consistent decrease of ECM biosynthetic tenocytes with DTR and aging, we also identified divergent retention of the remaining tenocytes between DTR and aged groups ***(Figure 3F-I)***. Broadly, tenocytes in the 6M DTR group express genes related ECM organization, development and remodeling such as *Edil3, Mmp2, Col5a1* and *Fn1*. Based on FEA, 6M DTR tenocytes are characterized as specialized cells for ECM organization and remodeling (“ECM organization”, “extracellular structure organization” “supramolecular fiber organization”, and “tissue remodeling”). In contrast, tenocytes from 21M B6 mice express genes related to hallmarks of aging like loss of proteostasis (*Ppp1r15a, Hspa8, Hsp90aa1, Hspa5, Hspb1, Hspa1, Hspa1b*, and *Dnajb9*) and inflammaging (*Il6, Stat3, Tnfaip3, Cxcl12*, and *Bcl3*). Indeed, FEA of 21M B6 tenocytes demonstrated processes related to “response to stress”, “response to unfolded protein”, “T cell and leukocyte activation”, “TNF and IL1 production”, and “aging” ***(Figures 3F, I)***. Further analysis showed that in the DTR group there was a retention of tenocytes 2, while this population was decreased by 96.2% in the aged WT tendon relative to the young WT **(*Figure 4B-E)***. Tenocytes 2 express genes that give rise to ECM proteins such as *Mmp2, Lum, Col1a1, Col3a1, Col11a1, and Mfap5*. FEA demonstrated biological processes such as “ECM and collagen fibril organization”, “supramolecular fiber organization”, “cell adhesion”, and “wound healing”, suggesting that tenocytes 2 exhibit functions related to ECM organization and wound healing ***(Figure 4F, H)***. Taken together, these divergent retention of tenocytes in DTR and aged tendons might dictate divergent healing responses after injury.

In addition to divergence of tenocyte subpopulations shifts, we also assessed for potential intrinsic programmatic shifts of tenocytes 1 and 2 with DTR and natural aging by comparing the transcriptomic profile of tenocytes 1 with DTR and aging ***(Figure 5A-J)***, as well as the transcriptomic profile of tenocytes 2 in the 6M WT vs 6M DTR groups ***(Figure 5K-O)***. In DTR compared to WT tendons, tenocytes 1 exhibited a total of 413 DEG ***(Figure 5A, B)***. FEA revealed that tenocytes 1 in the 6M WT group exhibited processes such as *“*translation”, “macromolecule biosynthetic process”, “structure morphogenesis”, while in the 6M DTR group, tenocytes 1 exhibited processes such as “cell migration”, “ECM organization”, and “wound healing” ***(Figure 5C-E)***. In contrast, looking at age-related programmatic shifts of tenocytes 1 ***(Figure 5F-H)***, while young WT tenocytes 1 exhibited processes such as “collagen biosynthesis”, “ECM organization”, and “immune cell chemotaxis” ***(Figure 5I)***, aged tenocytes 1 exhibited processes related to potential indicators of aging hallmarks such as loss of proteostasis “response to unfolded protein”, “protein refolding”, and ‘inflammaging’ “T cell activation”, “immune response” ***(Figure 5J)***. Regarding programmatic changes of tenocytes 2 in 6M WT vs DTR ***(Figure 5K)***, FEA analysis from the DEGs ***(Figure 5L, M)*** showed that tenocytes 2 in the young WT tendons exhibited biological processes such as *“macromolecule biosynthetic and metabolic processes”, “collagen metabolic processes”*, and *“ECM organization”*, while tenocytes 2 in the young DTR group exhibited further elevated as well as new processes related to matrix organization and remodeling such as *“ECM organization”, “extracellular fibril organization”*, and *“ECM assembly”* ***(Figure 5N, O)***. These divergent intrinsic programmatic shifts in tenocytes between DTR and aged tendons might also dictate the divergent healing responses that are observed in these models.

**Figure 5.**
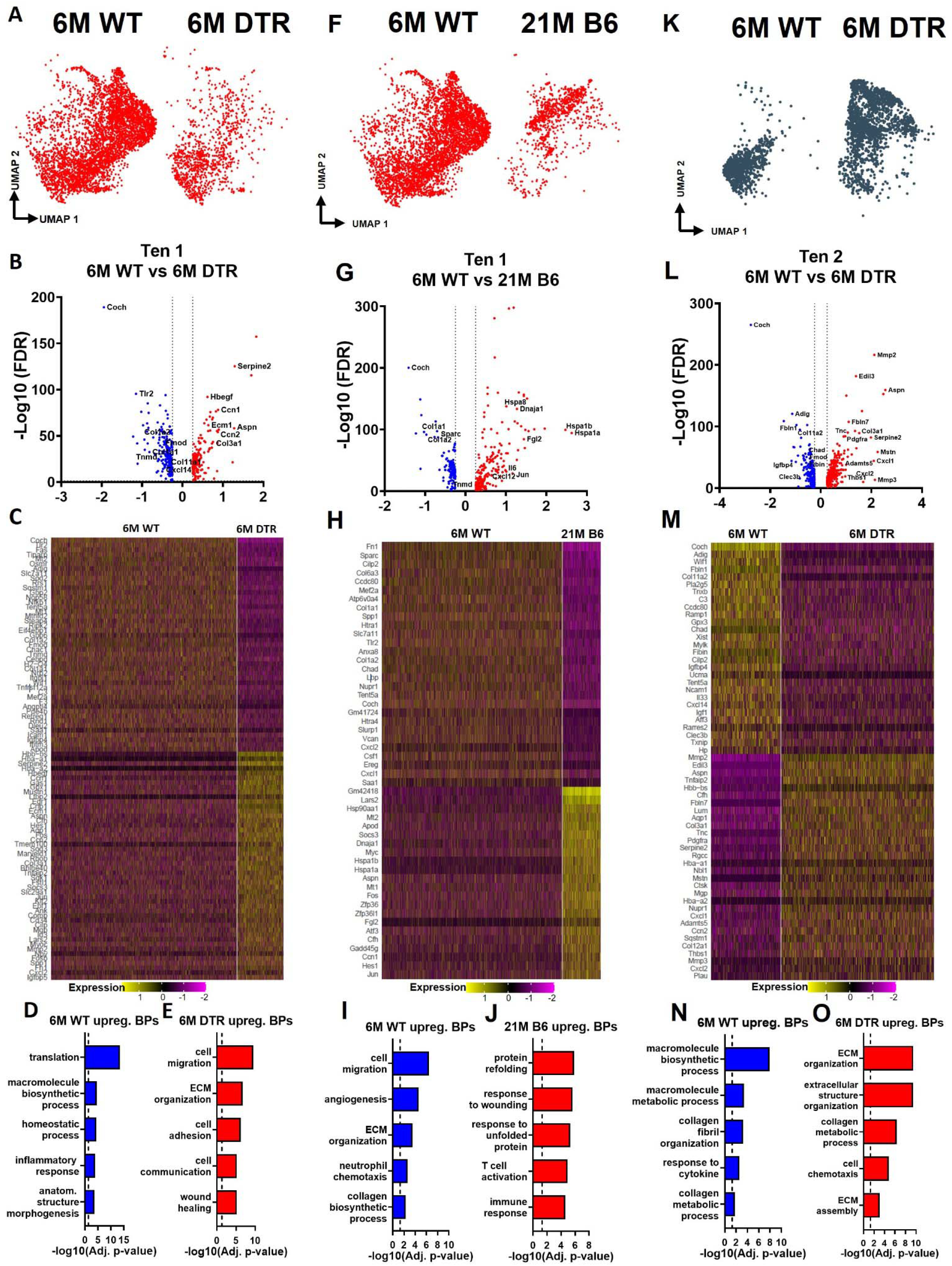
Tenocytes 1 and 2 become more ECM organizational in DTR tendons while tenocytes 1 exhibit indications of aging hallmarks such as loss of proteostasis and inflammaging. **(A)** UMAP plot of tenocytes 1 in 6M WT and DTR groups. **(B)** Volcano plot visualizing all the significantly different genes of tenocytes 1 between 6M WT (left side) and 6M DTR (right side). **(C)** Heatmap of the top 50 differentially expressed genes of tenocytes 1 between 6M WT and DTR groups. **(D)** Biological processes of tenocytes 1 in 6M WT group. **(E)** Biological processes of tenocytes 1 in 6M DTR group. **(F)** UMAP plot of tenocytes 2 in 6M WT and 6M DTR groups. **(G)** Volcano plot visualizing all the significantly different genes of tenocytes 2 between 6M WT (left side) and 6M DTR (right side). **(H)** Heatmap of the top 50 differentially expressed genes of tenocytes 2 between 6M WT and 6M DTR groups. **(I)** Biological processes of tenocytes 2 in 6M WT group. **(J)** Biological processes of tenocytes 2 in 6M DTR group. **(K)** UMAP plot of tenocytes 1 in 6M and 21M B6 groups. **(L)** Volcano plot visualizing all the significantly different genes of tenocytes 1 between 6M WT (left side) and 21M B6 (right side) groups. **(M)** Heatmap of the top 50 differentially expressed genes of tenocytes 1 between 6M and 21M B6 groups. **(N)** Biological processes of tenocytes 1 in 6M WT group. **(O)** Biological processes of tenocytes 1 in 21M B6 group.

### Tenocyte autocrine and paracrine signaling is differentially altered with *Scx*^*Lin*^ depletion vs. natural aging

Cellular communication is crucial for maintenance of tissue homeostasis [39-43], and physiological communication patterns are impaired with aging, leading to disruptions tissue health [44, 45]. Therefore, we next determined what the predominant tenocyte communication patterns are in young WT tendon, and how these patterns are changed in the contexts of depletion and natural aging.

Young WT tenocytes 1-3 exhibit both autocrine and paracrine signaling ***(Fig. S6A, B)***. Tenocytes 1 exhibit high communication strength as both sender and receiver, tenocytes 2 is the predominant sender but exhibits low intercellular communication strength as a receiver and tenocytes 3 exhibits high communication strength as a receiver but low strength as a sender ***(Fig. S6C, D)***. *Scx*^*Lin*^ cell depletion resulted in substantial shifts in tenocyte autocrine and paracrine communication ***(Figure 6A-F)***. Tenocytes 1 exhibited a decrease in overall communication, tenocytes 2 exhibited an increase in autocrine and paracrine signaling to tenocytes 1, but a decrease in paracrine signaling to tenocytes 3. Finally, tenocytes 3 demonstrated a decrease in autocrine and paracrine signaling to tenocytes 1, and an increase in paracrine signaling to tenocytes 2 ***(Figure 6B)***. Five signaling pathways (CHAD, LIFR, NECTIN, CADM, EPHA) were identified exclusively in the 6M WT group, while four signaling pathways (IL6, SEMA3, NOTCH, and FGF) were expressed only in the 6M DTR group ***(Figure 6C)***. The heatmaps demonstrate the strength of outgoing (***Fig. 6D***), and incoming (***Fig. 6E***) signaling patterns in tenocytes. We further focused on the THBS signaling pathway since *Thbs2, Thbs3*, and *Comp* were significantly decreased with depletion in both the proteomics and scRNAseq datasets ***(Figure 2A, D; Fig. S5)***. More specifically, we identified that *Thbs3-Sdc1, Thbs2-Sdc4, Thbs2-Sdc1*, and *Thbs2-Cd47* ligand-receptor interactions were decreased in tenocytes 1-3 with depletion ***(Figure 6F)***, suggesting that this interaction may be important for maintenance of tendon homeostasis.

**Figure 6.**
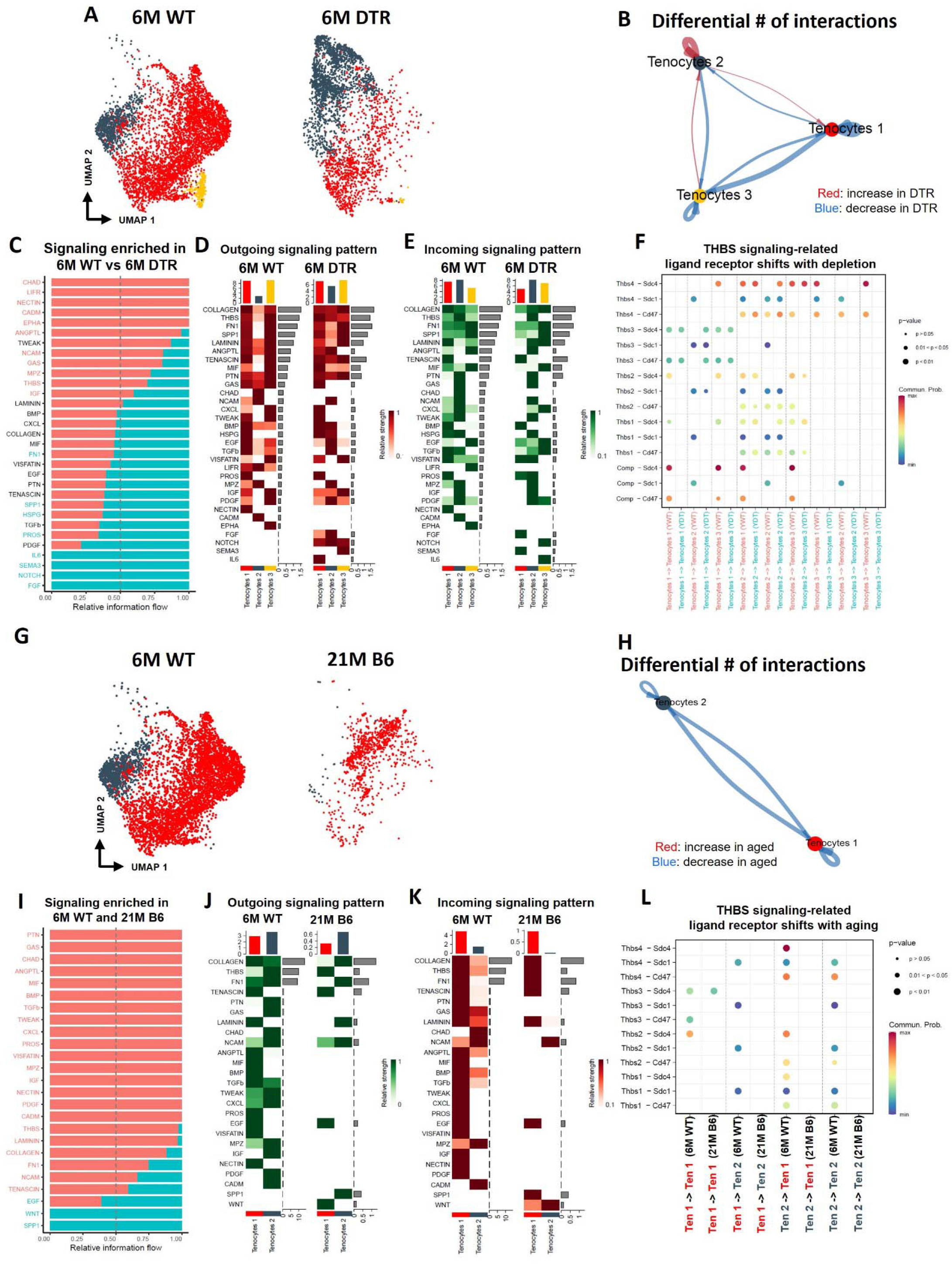
Tenocyte-tenocyte communication is impaired with both *Scx*^*Lin*^ cell depletion and natural aging. **(A)** UMAP plot of tenocytes subpopulations in the 6M WT and DTR groups. **(B)** Differential number of cell-cell interactions of tenocytes 1-3 in the 6M WT and DTR groups (red color indicates an increase and blue color indicates a decrease in cell communication with *Scx*^*Lin*^ cell depletion relative to WT, respectively; arrow indicates the communication between two different cells and the direction of the arrow indicates the cell that expresses the ligand vs the cell that expresses the receptor, where the pointed cell by the arrow is the receiver). **(C)** All the signaling pathways that were identified in the 6M WT and DTR groups (pathways highlighted with orange color indicate those that were expressed only or higher in the 6M WT relative to 6M DTR group; pathways highlighted with green color indicate those that were expressed only or higher in the 6M DTR relative to 6M WT group). **(D)** Heatmap with all the outgoing signaling patterns for tenocytes 1-3 for the 6M WT and DTR groups. **(E)** Heatmap with all incoming signaling patterns for tenocytes 1-3 for the 6M WT and DTR groups. **(F)** THBS signaling-related ligand-receptor interactions that take place in the 6M WT and DTR groups. **(G)** UMAP plot of tenocytes subpopulations in the 6M WT and 21M B6 groups. **(H)** Differential number of cell-cell interactions of tenocytes 1 and 2 in the 6M WT and 21M B6 groups (blue color indicates a decrease in cell communication with natural aging relative to WT; arrow indicates the communication between two different cells and the direction of the arrow indicates the cell that expresses the ligand vs the cell that expresses the receptor, where the pointed cell by the arrow is the cell that expresses the receptor or in other words the receiver **(I)** All the signaling pathways that were identified in the 6M WT and 21M B6 groups (pathways highlighted with orange color indicate those that were expressed only or higher in the 6M WT relative to the 21M B6 group; pathways highlighted with green color indicate those that were expressed only or higher in the 21M B6 groups relative to 6M WT group). **(J)** Heatmap with all the outgoing signaling patterns for tenocytes 1 and 2 for the 6M WT and 21M B6 groups. **(K)** Heatmap with all incoming signaling patterns for tenocytes 1 and 2 for the 6M WT and 21M B6 groups. **(L)** THBS signaling-related ligand-receptor interactions that take place in the 6M WT and 21M B6 groups.

**Figure 7.**
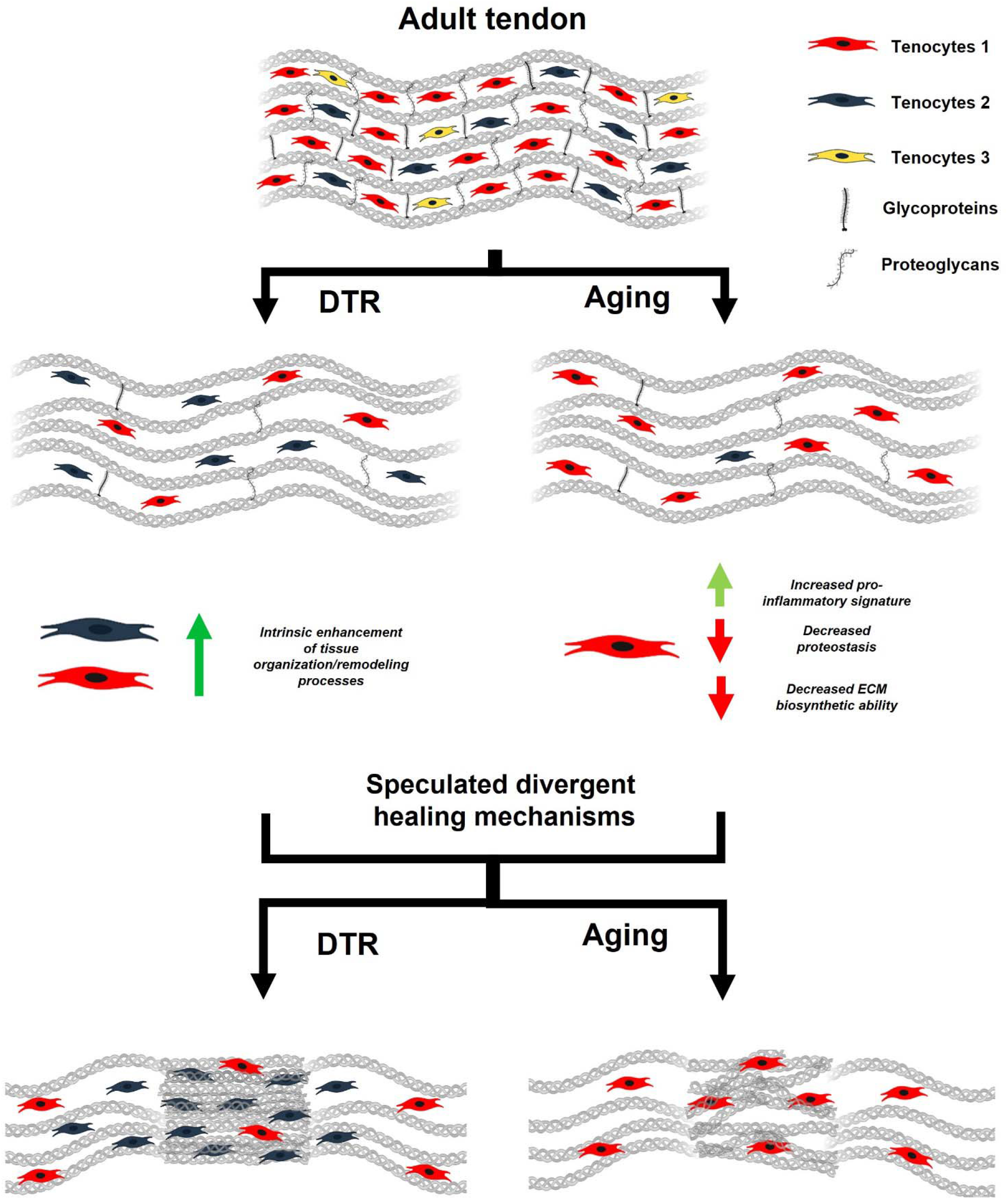
Schematic highlighting key findings and proposed models for divergent healing responses in young depleted vs aged tendons. Both young DTR and aged WT tendons have similar decreases in tissue structure, organization, material quality, and total cell density. They also follow the same mechanism of ECM degeneration via a substantial decrease in the number of proteoglycans and glycoproteins with high turnover rate. However, in terms of cell composition shifts with depletion and natural aging, young DTR tendons are comprised mainly of tenocytes specialized in tissue organization and remodeling. What is more, young DTR have little to no inflammatory/reactive tenocytes. In contrast, old WT tendons are comprised predominantly of inflammatory and age-related functionally impaired tenocytes and exhibit little to no presence of tissue remodeling tenocytes. We speculate that these significantly different cellular compositions are some of the main causes resulting in divergent healing responses, with young DTR tendons showing improved healing while old WT tendons exhibiting impaired healing response.

In contrast to altered communication in response to DTR, natural aging results in an almost complete loss in tenocyte communication compared to 6M WT ***(Figure 6G-L)***. Tenocytes 1 and 2 exhibited a loss of both autocrine and paracrine communication ***(Figure 6H)***. A majority of the signaling pathways were expressed only in the 6M WT group, suggesting that with natural aging there is a substantial and widespread loss of tenocyte-tenocyte communication ***(Figure 6I-K)***. Interestingly, SPP1 and WNT pathways were present only in the aged group in both an outgoing and incoming pattern ***(Figure 6J, K)***. Finally, we also identified a loss of THBS signaling with natural aging, including *Thbs3-Sdc4, Thbs3-Sdc1, Thbs2-Sdc4, Thbs2-Sdc1*, and *Thbs2-Cd47* ***(Figure 6L)***, which was very similar to the ligand-receptor interactions lost with depletion ***(Figure 6F)***. Taken together, aging results in broad decline in tenocyte-tenocyte communication, and future work will define whether decreased and altered communication between the remaining tenocytes may drive further degeneration beyond that observed at 21M B6.

## Discussion

Aging impairs tendon homeostasis, increases the risk of degeneration [24, 46] and the frequency of injury, and diminishes healing capacity after injury [38, 46, 47]. In addition, clinical studies have shown that age-related tendon changes are associated with impaired dynamic stability and increased risk of falling, factors that significantly increase both morbidity and mortality [48-50]. In this study, we identified the cell and molecular mechanisms that maintain tendon homeostasis during adulthood and demonstrate how these mechanisms are disrupted with aging. Moreover, we demonstrated that depletion of *Scx*^*Lin*^ cells in the adult tendon mimics age-related impairments in cellular density, ECM structure, composition, and material quality. By comparing and contrasting these two models we were able to delineate the mechanisms that regulate tendon homeostasis, independent of the systemic effects of aging, while also establishing the tendon-specific response to aging. More specifically, we have demonstrated that loss of a critical mass of biosynthetic tenocytes, or biosynthetic tenocyte function initiates a degenerative cascade in tendon. This is particularly striking as it demonstrates that it is the decline in cell density and the accompanying loss of cell function itself that drives age-related tendon degeneration, rather than age-related programmatic shifts. Finally, these data suggest that the initiation and early evidence of degeneration occurs much earlier in life than previously appreciated. As such, we have identified prevention of biosynthetic tenocyte apoptosis as a critical intervention point for maintaining tendon structure-function.

While we have previously shown that this *Scx*^*Lin*^ depletion strategy results in a ∼60% decrease in tendon cell density [36], it was unknown whether this was a sustained or transient depletion event that eventually results in re-population of the tendon via compensation by non-depleted populations. Indeed, this model results in sustained cell depletion, thus allowing assessment of the long-term impact of *Scx*^*Lin*^ cell deficiency. It is important to note that this model likely depletes any “progenitor” cell populations due to the use of non-inducible Scx-Cre, further inhibiting any potential cellular rebounding post-depletion. We observed that *Scx*^*Lin*^ cell depletion resulted in almost identical cellular density with older (12M old) and geriatric (31M old) tendons, indicating that with depletion, there is an acceleration of the natural age-related tendon cell loss. An age-related decline in tendon cell density has been extensively documented, with consistent decreases across anatomically distinct tendons and animal models [51-55]. However, the fact that Scx^Lin^ depletion in young animals results in a comparable decline in cellularity suggests there may be mechanisms that maintain a minimum level of cellularity and prevents complete cell loss, although these mechanisms remain to be defined. Moreover, the recapitulation of many established hallmarks of tendon aging, including altered ECM structure, organization, and material quality [17, 19, 25, 52, 56-58], further supports the utility of defining the consistent underlying mechanisms between DTR and aging to define previously unappreciated drivers of age-related tendon pathology. Indeed, the combination of transcriptomic and proteomic analyses in these models has facilitated identification of the key alterations in ECM composition that are associated with initiation and progression of tendon degeneration. Consistent with declines in high-turnover rate proteoglycans and glycoproteins with aging in other tissues [10, 59-61], these classes of proteins were decreased in both aged and young depleted tendons, demonstrating that *Scx*^*Lin*^ cells are required to directly regulate the synthesis and maintenance of multiple GPs and PGs, which are crucial to maintain tendon structure-function, since their decrease results in significantly impaired collagen fibril organization and biomechanical properties. While other studies have shown through genetic KO models, that high turnover rate GPs and PGs are required to maintain tissue (tendon, bone, cartilage, skin) homeostasis during adulthood [57, 58, 62-73], they have been primarily descriptive. In contrast, we demonstrate that *Scx*^*Lin*^ cells are a “master-regulator” of multiple ECM-related proteins during tendon homeostasis. More specifically, we identified decreased expression of COCH and CHAD protein in the tendon ECM in these models, as well as a consistent decline in *Coch*+ and *Chad*+ cells. Interestingly, COCH expression is observed in the spiral ligament of the inner ear, and *Coch*-/- results in degeneration of this ligament [74], further supporting the potential importance of this molecule in maintenance of tendon structure.

Consistent with prior scRNAseq studies in mouse and human tendons [2, 4, 75], we have identified substantial heterogeneity of the tendon cell environment. We have further built on this by establishing the functional consequences of decreasing the subpopulation that is primarily characterized by biosynthetic functions. Interestingly, age-related declines in ECM biosynthetic function have also been identified in mouse and human skin fibroblasts [76, 77], thus, future work will focus on establishing and disrupting the mechanisms that drive age-related declines in biosynthetic tenocyte number and function.

In addition to comparable declines in biosynthetic function and subsequent initiation of tendon degeneration in these models, there are several informative areas of divergence. First, the tenocytes that remain in aged tendons demonstrate programmatic skewing, including an increased pro-inflammatory signature and loss of proteostasis. While these shifts are not necessary for homeostatic disruption, as evidenced by comparable disruptions in DTR tendons in the absence of this skewing, it is possible that these changes drive subsequent degeneration at timepoints beyond those included in this study. Consistent with this, 31M B6 tendons exhibit substantial increases in ECM disorganization, relative to DTR. Moreover, while no enrichment of senescent markers was observed in aged tenocytes, it is possible that the pro-inflammatory signature and loss of proteostasis may be a precursor to subsequent tenocyte senescence, which could be an important driver of additional tendon degeneration. Another intriguing consideration is how these divergent shifts in tenocyte programs may relate to the divergent healing outcomes observed between these models. Consistent with impaired healing capacity in many aged tissues [38, 78-80], we have previously shown that 22M tendons heal in a mechanically inferior manner, relative to young tendons, and fail to form the provisional ECM needed for successful healing [38]. Thus, establishing how these intrinsic programmatic shifts may impair specific aspects of the healing process will be necessary to develop strategies to rescue the aged tendon healing process. In contrast, the cells that remain in DTR tendons display programmatic shifts associated with enhanced remodeling capacity. Moreover, DTR tendons demonstrate improved healing capacity relative to age-matched WT repairs [36]. As such, understanding the signals that drive this programmatic shift in DTR, and establishing the mechanisms by which Scx^Lin^ depletion enhances tendon healing holds tremendous potential for tendon tissue engineering and therapeutic strategies. Finally, these data highlight the surprising finding that homeostatic disruptions including decreased cell density and altered ECM composition, quality and organization are not of central relevance to the quality of the tendon healing response, suggesting a de-coupling of the mechanisms that regulate homeostasis and healing capacity.

One limitation of the study is that the *Scx-Cre; DTR* ^*F/+*^ targets all *Scx*^*Lin*^ cells, and the *Scx*^*Lin*^ is known to contribute to tissues in addition to tendon (*e*.*g*. bones and muscle) [81, 82]. However, our initial plans to deplete *Scx*^*Lin*^ cells by utilizing the inducible *Scx-Cre*^*ERT2*^ crossed to the diphtheria toxin A-subunit gene (DTA) mouse model, resulted in insufficient cell depletion. *Scx-Cre*^*ERT2*^; Rosa-DTR mice could be an alternative to inducibly deplete only adult *Scx*^*Lin*^ cells, although this model requires both TMX and DT administration, and discrepencies in labelling vs. depletion may be observed. In addition, while we clearly demonstrate programmatic skewing in aged (21M) tendons, relative to young, it is not clear when cells begin to undergo this programmatic shift. Future work to define the temporal nature will the establish the onset of these shifts as an additional important intervention point for maintaining physiological tendon cell function.

In summary, our morphological and multi-omics analyses identified Scx^Lin^ cell depletion as a novel model of accelerated tendon ECM aging based on comparable changes in ECM organization, composition, and material quality with aged tendons. Moreover, we defined loss of a critical mass of biosynthetic tenocyte function as a conserved initiator of disrupted tendon homeostasis and degeneration in both young and aged animals. Thus, identifying strategies to prevent apoptosis of this population is an important priority for maintaining tendon health through the lifespan. Finally, we have demonstrated divergent programmatic skewing between Scx^Lin^ cell depletion in young animals, and natural aging, and suggest that these differences may underpin the divergent healing capacity observed in these models. Therefore, it will be necessary to develop different strategies to address or prevent these two distinct manifestations of aging in the tendon to preserve both tendon homeostasis and healing capacity.

## Author contributions

Study conception and design: AK, AEL; Acquisition of data: AK; Analysis and interpretation of data: AK, AECN, MRB, AEL; Drafting of manuscript: AK, AEL; Revision and approval of manuscript: AK, AECN, MRB, AEL.

## Data and material availability

All data needed to evaluate the conclusions in the paper are present in the paper or the Supplementary Materials.

## SUPPLEMENTARY FIGURES

**Supplement Figure 1.**
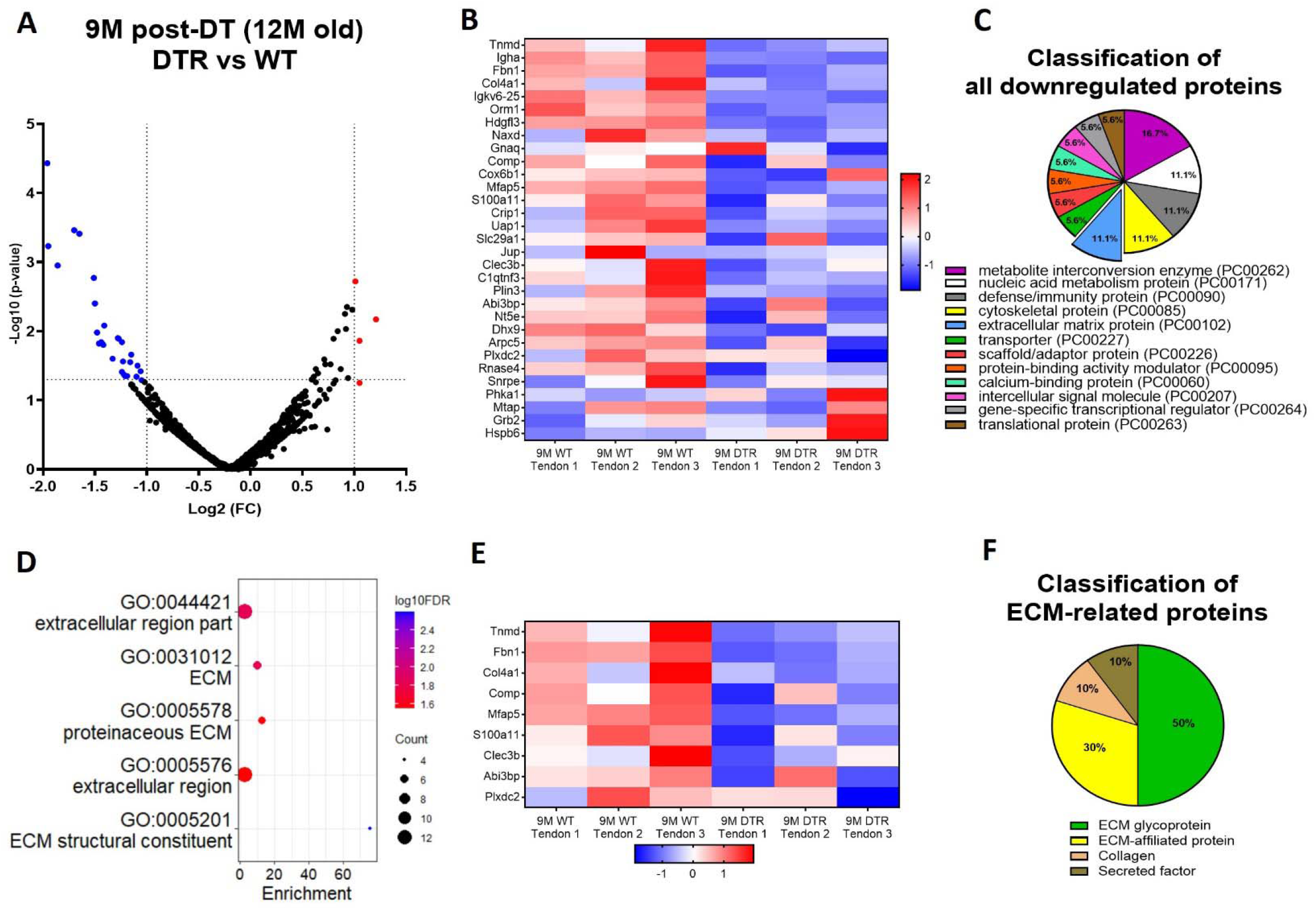
*Scx*^*Lin*^ cells maintain long term tendon homeostasis via synthesis of high turnover rate GPs and PGs. **(A)** Volcano plot and **(B)** heatmap visualizing the significantly different protein abundances between DTR and WT groups at 9M post-depletion. **(C)** Classification of all downregulated proteins between the DTR and WT FDL tendons at 9M post-depletion. **(D)** Functional enrichment analysis of cellular components and molecular functions of all downregulated proteins between the DTR and WT FDL tendons at 9M post-depletion. **(E)** Heatmap of all differentially abundant ECM-related proteins between the DTR and WT FDL tendons. **(F)** Classification of all ECM-related downregulated proteins between the DTR and WT FDL tendons.

**Supplement Figure 2.**
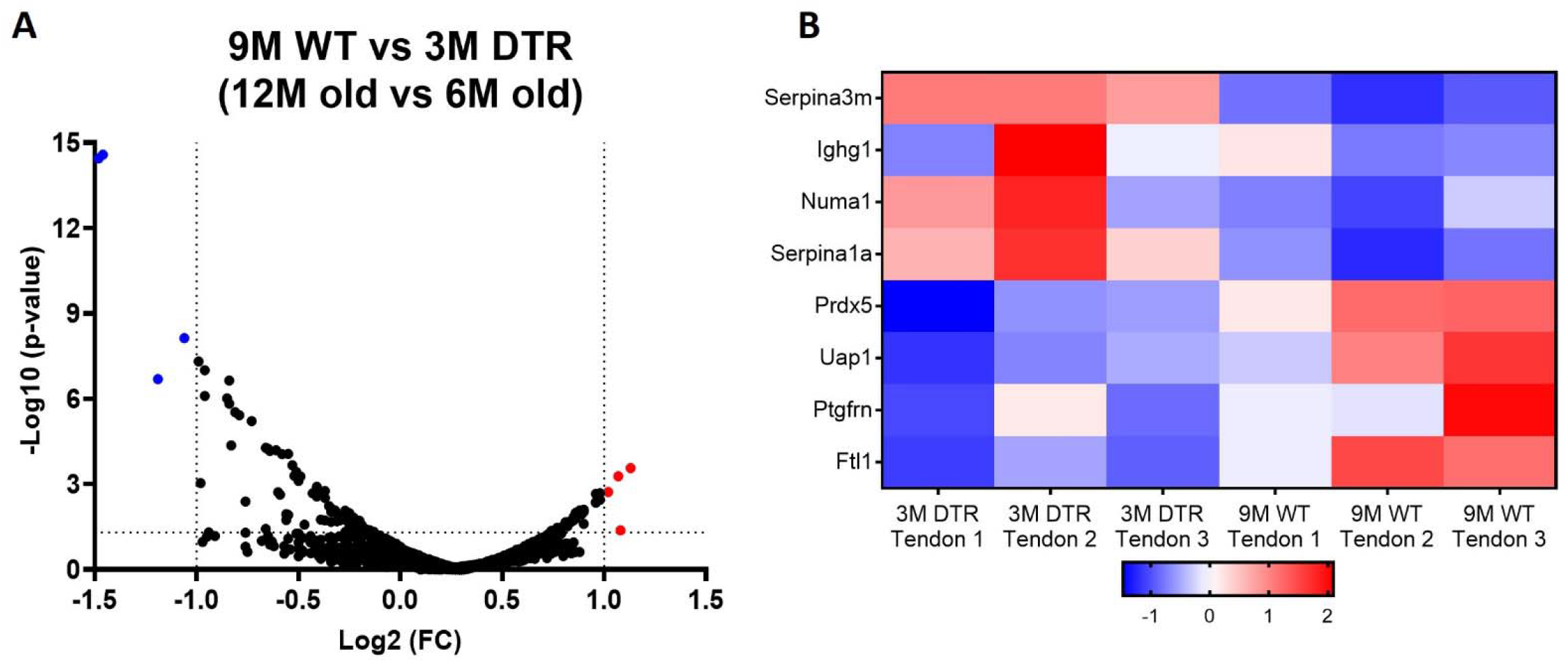
The proteome between old WT and young DTR tendons is almost identical with no differences in high turnover rate GPs and PGs. **(A)** Volcano plot and **(B)** heatmap visualizing the significantly different protein abundances between 12M old WT and 6M old DTR groups

**Supplement Figure 3.**
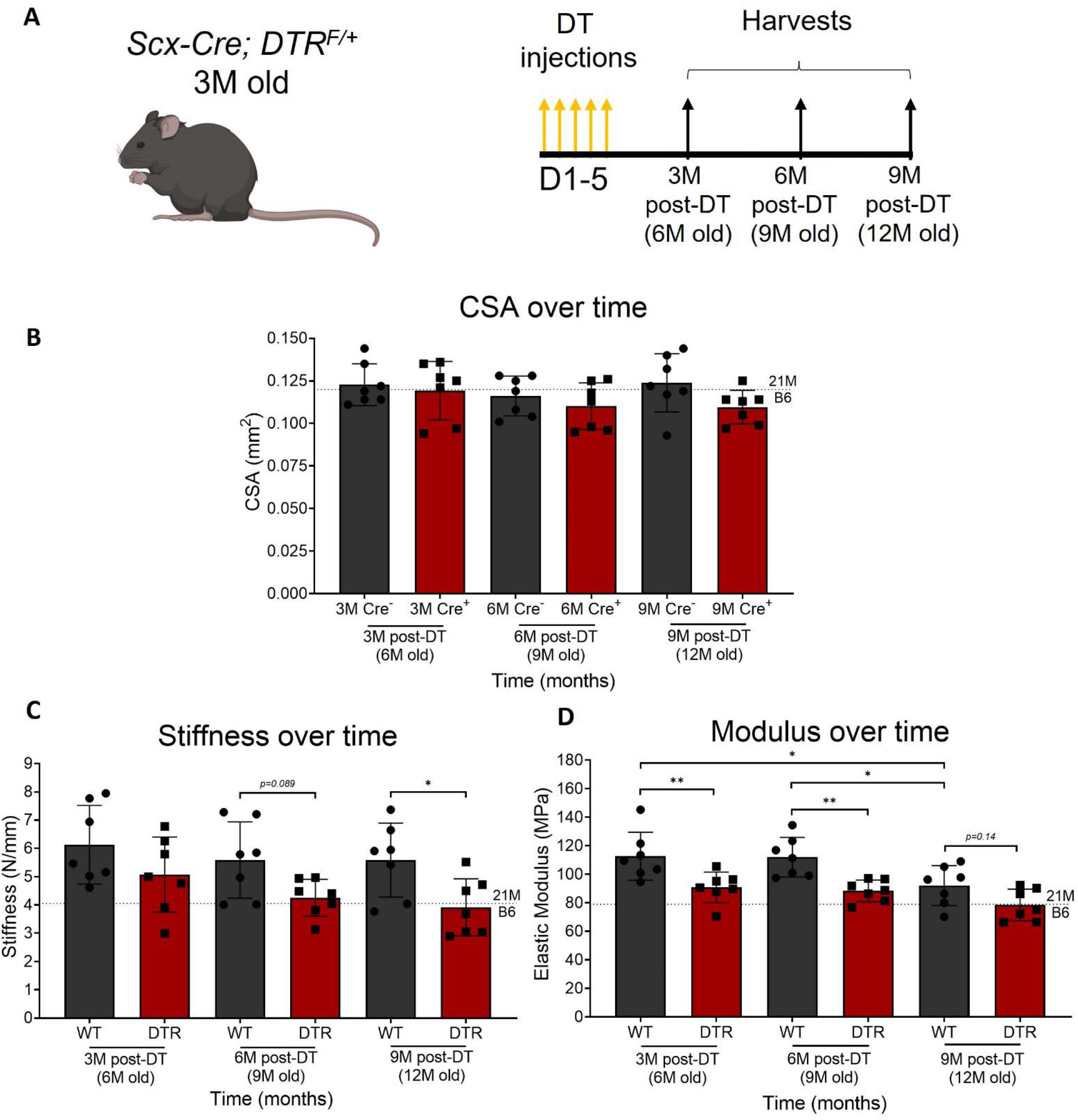
Depletion of *Scx*^*Lin*^ cells significantly impairs tendon structural and material properties. Timeline of DT injections and tissue harvesting **(A)**. Quantification of CSA **(B)**, stiffness **(C)**, and elastic modulus **(D)** of WT and DTR tendons at 3, 6, and 9 months post-depletion. CSA, stiffness, and elastic modulus between genotype (WT or DTR) and timepoint (3M, 6M, or 9M). N=3-5 per genotype. Two-way ANOVA with Sidak’s multiple comparisons test used to assess statistical significance of CSA, stiffness, and elastic modulus, * indicates p<0.05; ** indicates p<0.01; *** indicates p<0.001; **** indicates p<0.0001.

**Supplement Figure 4.**
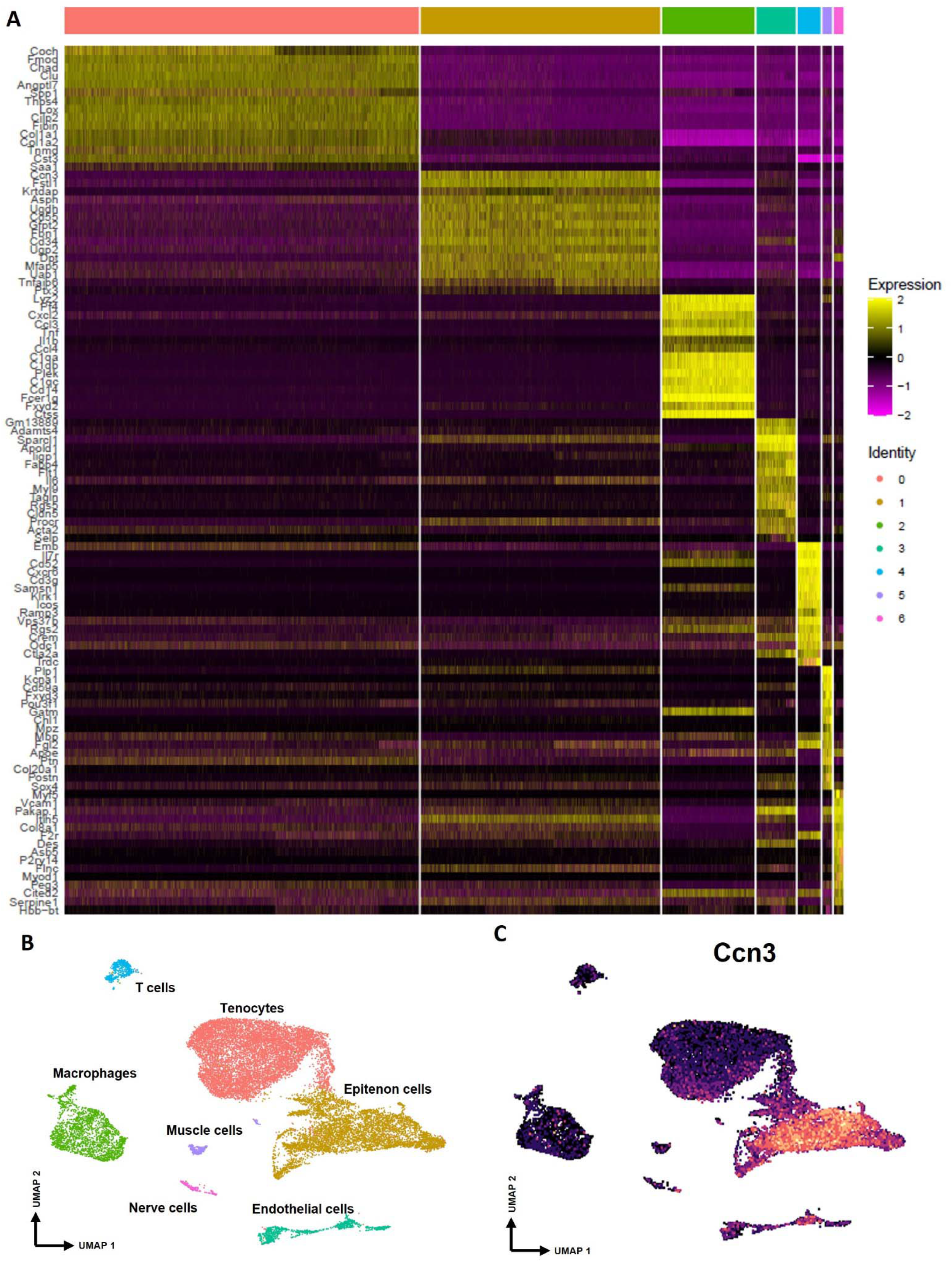
Annotation of scRNAseq-based identified cell clusters in the integrated data. **(A)** Heatmap of top 15 DEGs for each cluster. **(B)** UMAP with all different clusters annotated based on **(A). (C)** Feature plot of the epitenon marker *Ccn3* being expressed by the epitenon sub-cluster.

**Supplement Figure 5.**
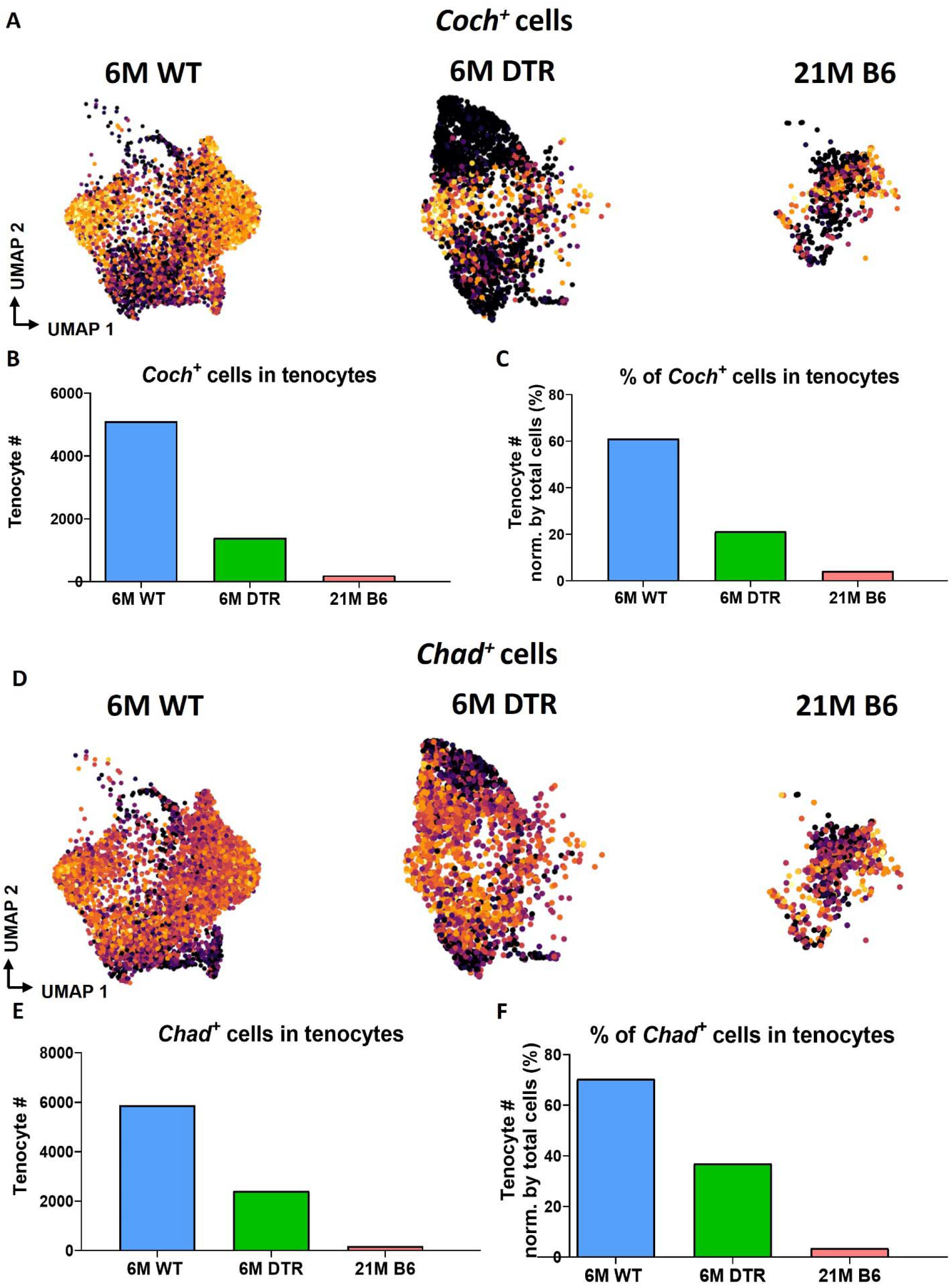
Significant decrease of *Coch*^*+*^ and *Chad*^*+*^ cells with *Scx*^*Lin*^ cell depletion and natural aging. **(A)** UMAP plots of *Coch*^*+*^ tenocytes in the 6M WT, DTR, and 21M B6 groups. **(B)** Total tenocyte number of *Coch*^*+*^ cells in each condition. **(C)** % of *Coch*^*+*^ normalized by total tenocytes for each condition. **(D)** UMAP plots of *Coch*^*+*^ tenocytes in the 6M WT, DTR, and 21M B6 groups. **(E)** Total tenocyte number of *Chad*^*+*^ cells in each condition. **(F)** % of *Chad*^*+*^ normalized by total tenocytes for each condition.

**Supplement Figure 6.**
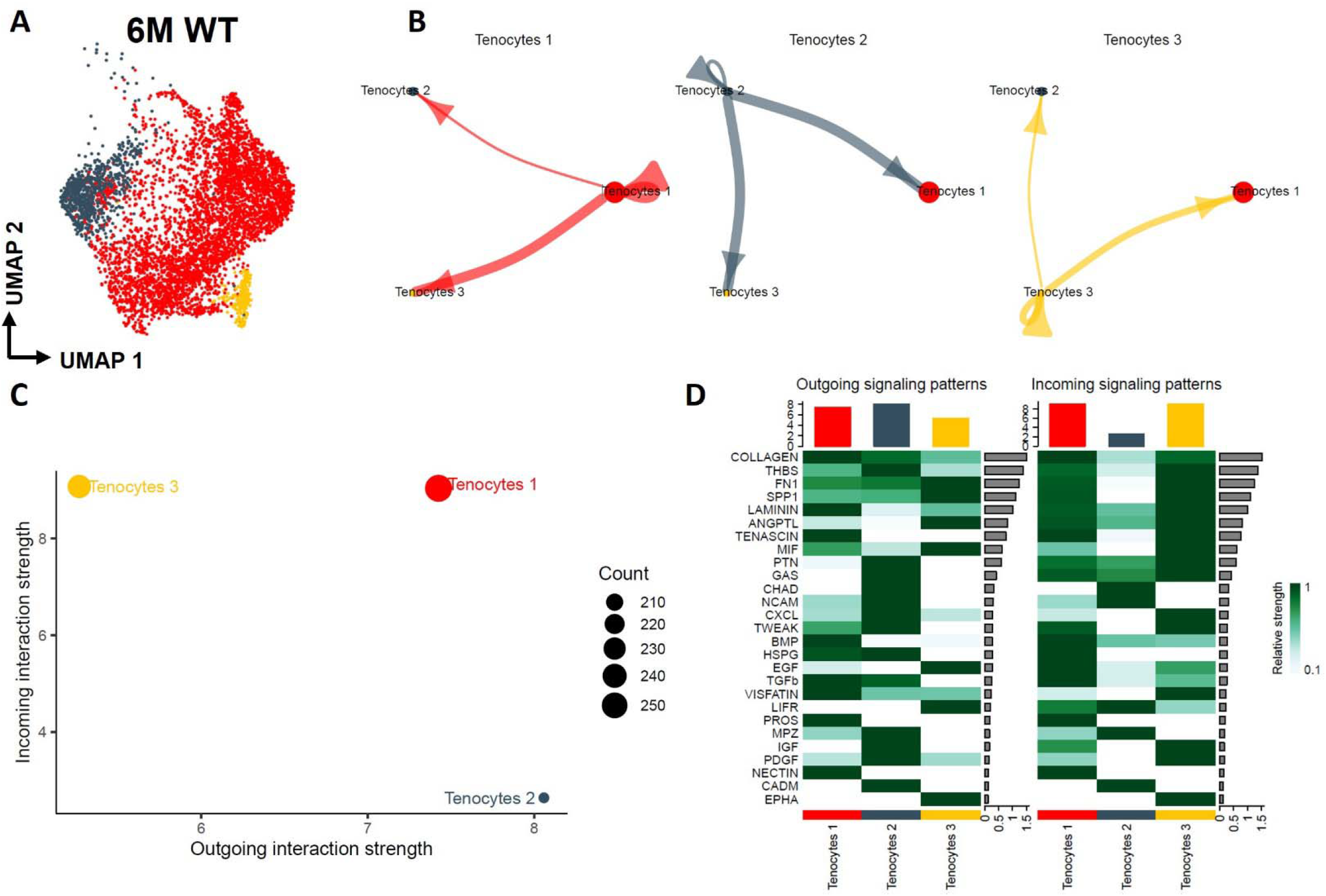
Tenocyte-tenocyte communication in the young adult FDL tendons. **(A)** UMAP plot visualizing the three tenocyte subpopulations that exist in the 6M WT FDL tendons. **(B)** Circle plot visualizing the autocrine and paracrine communication strength of each tenocyte subpopulation. **(C)** CellChat-based plot visualizing the incoming and outgoing communication strengths for each tenocyte subpopulation. **(D)** Heatmap of all the signaling pathways that are present during tenocyte-tenocyte communication, highlighting the outgoing vs incoming patterns for each tenocyte subpopulation.

**Supplement Figure 7.**
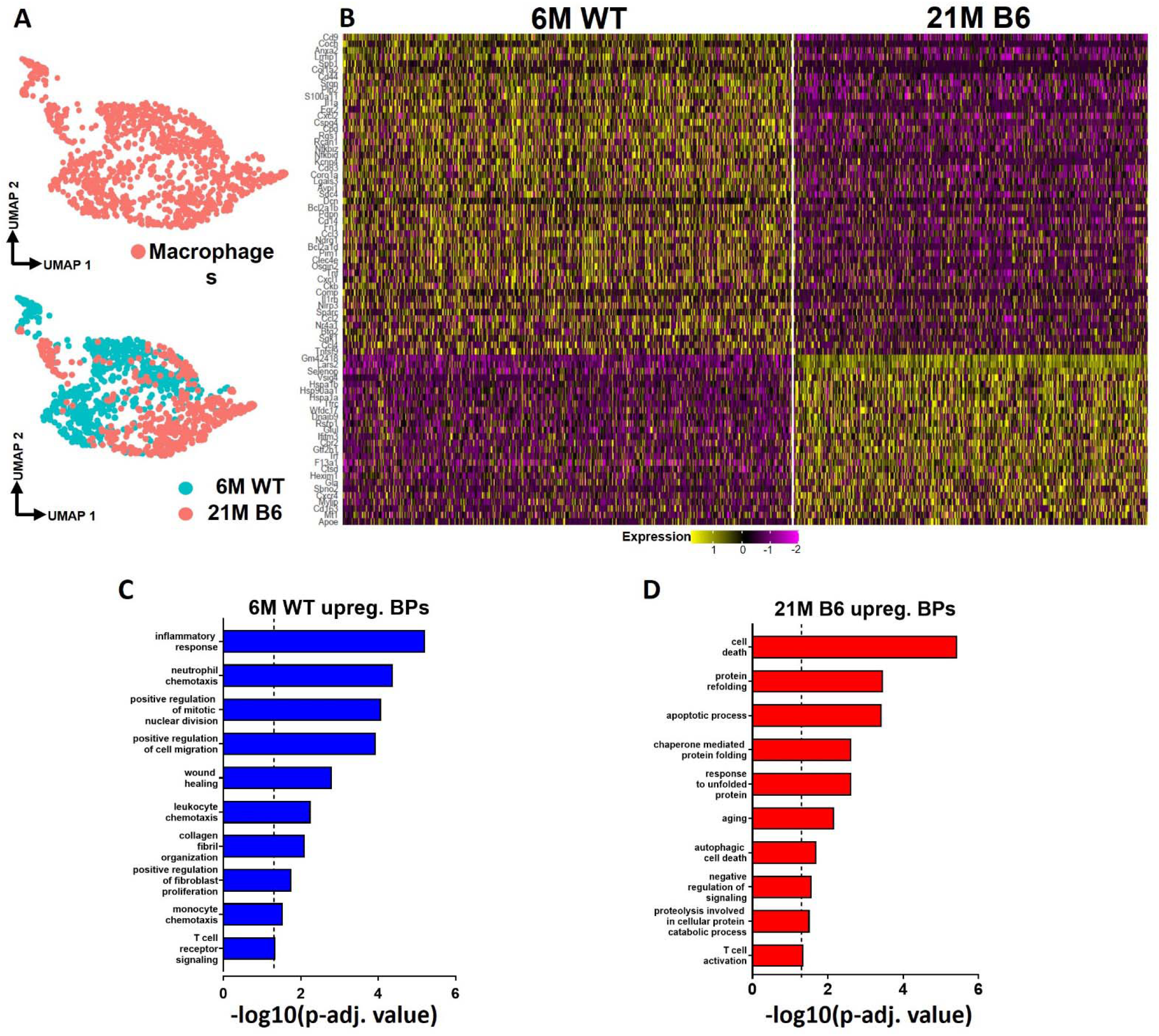
Tendon resident macrophages exhibit significant age-related intrinsic programmatic shifts. **(A)** UMAP plot of the overall tendon resident macrophages as well as separated based on age (6M WT and 21M old B6). **(B)** Heatmap with the top 50 DEG of tendon resident macrophages between the young vs old WT groups. **(C)** BPs enriched in the young WT resident tendon macrophages based on the DEGs. **(D) (C)** BPs enriched in the aged WT resident tendon macrophages based on the DEGs.

