## Supplementary figures and images for "Scleraxis-lineage cells are required for tendon homeostasis and their depletion induces an accelerated extracellular matrix aging phenotype"

### Supplemental figure 1

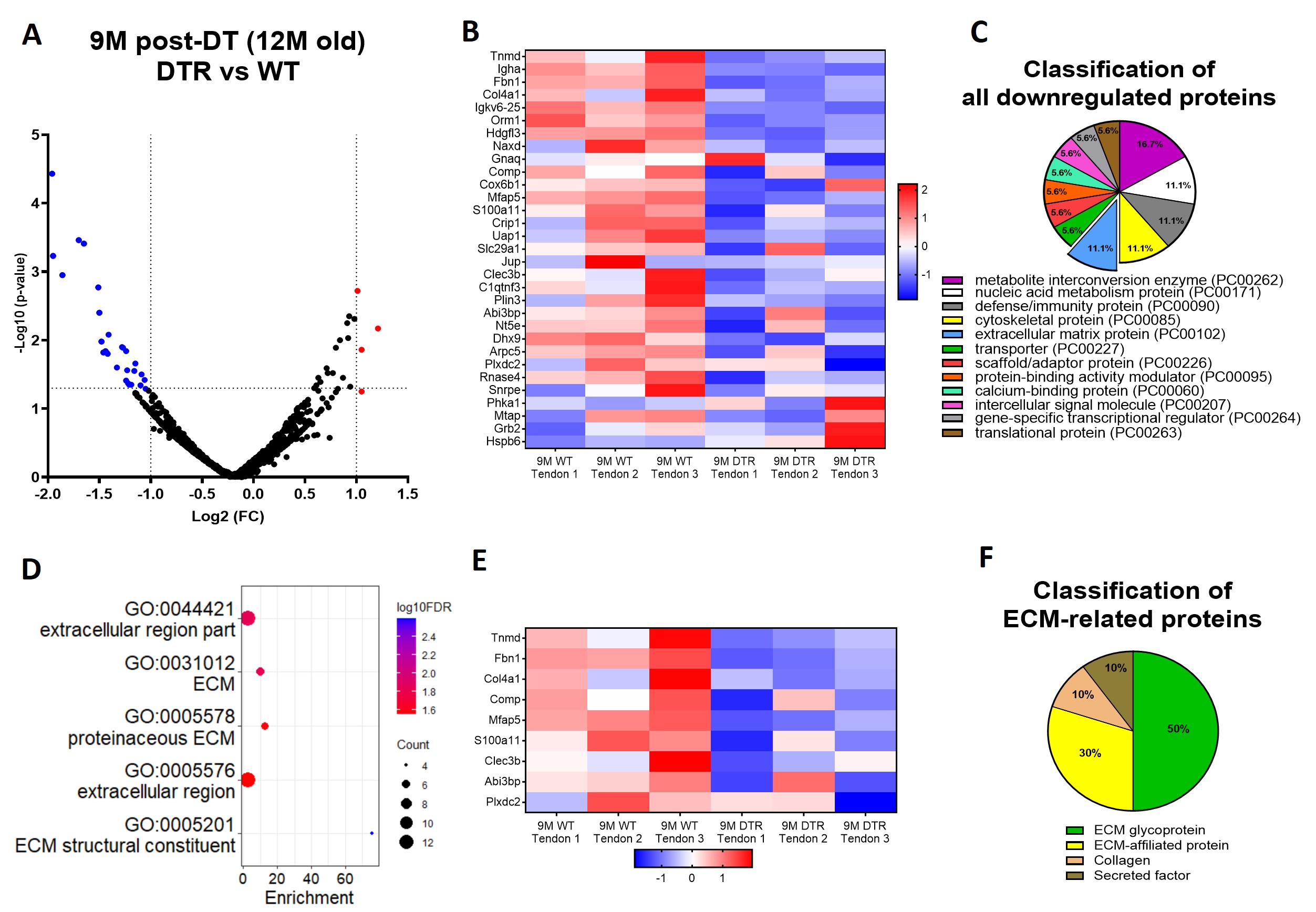

### Supplemental figure 2

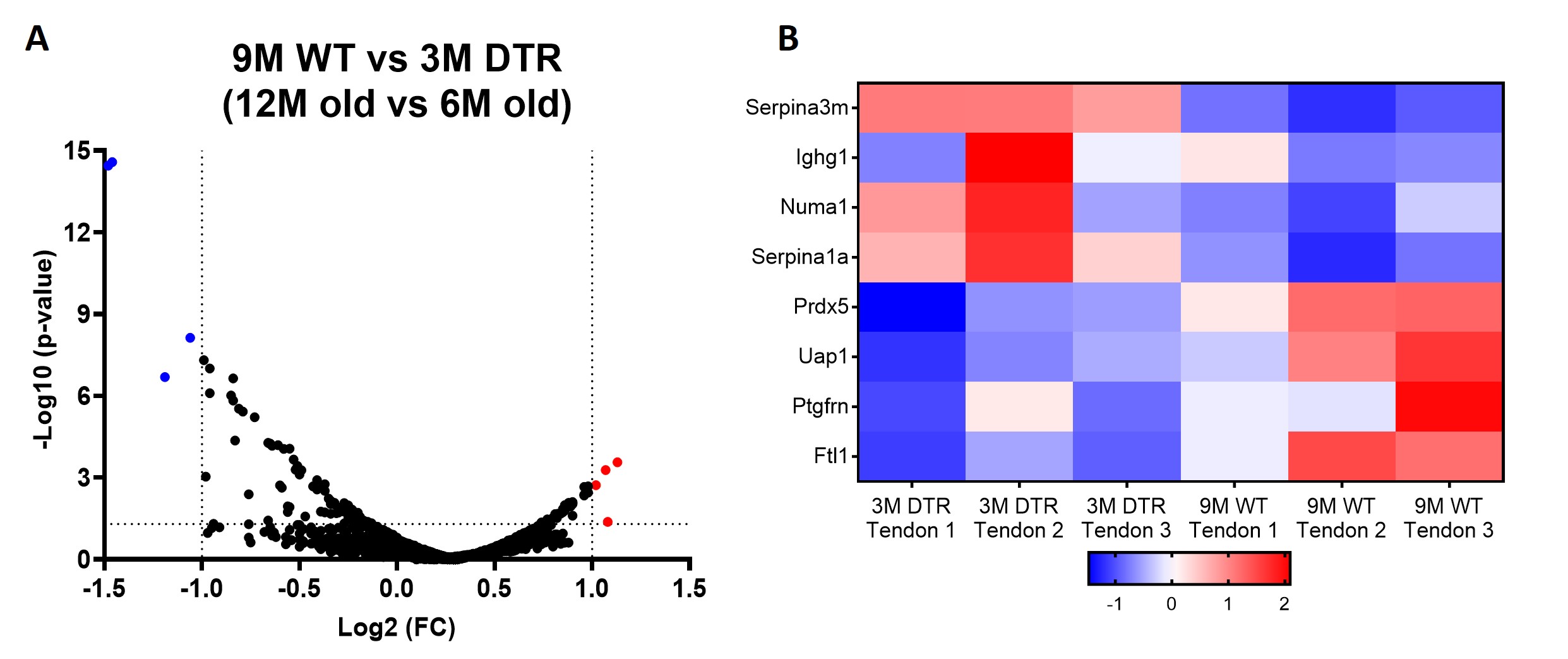

### Supplemental figure 3

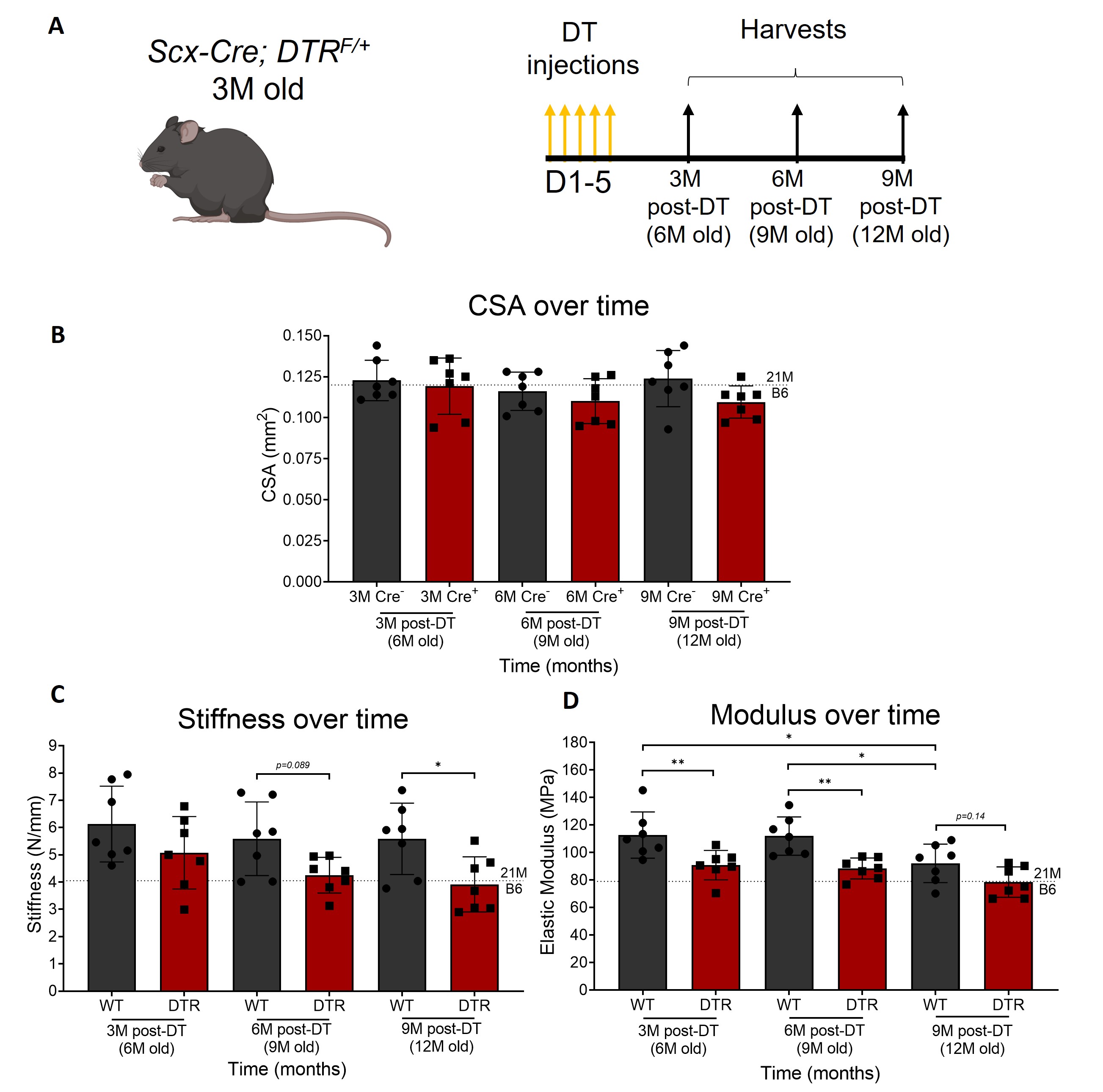

### Supplemental figure 4

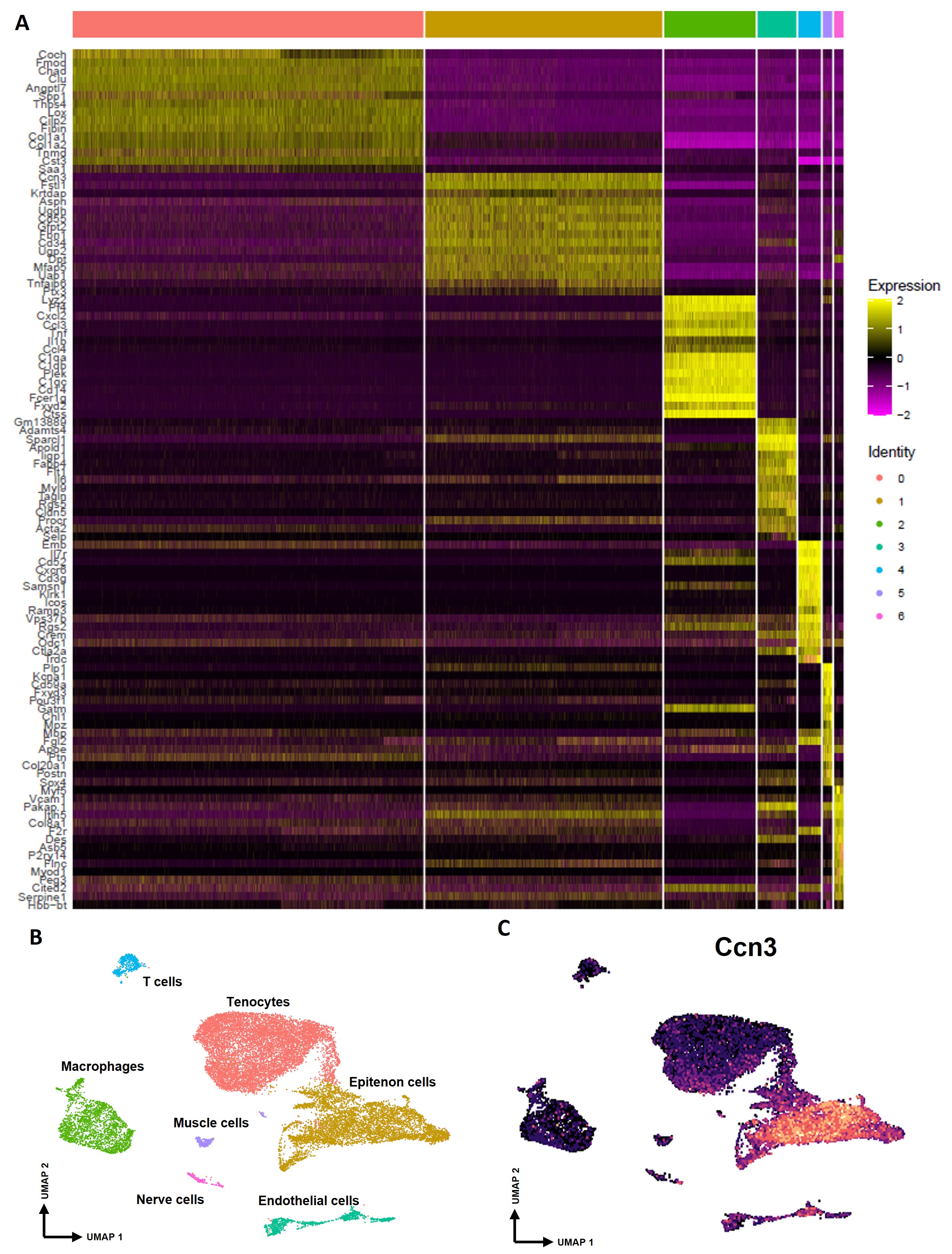

### Supplemental figure 5

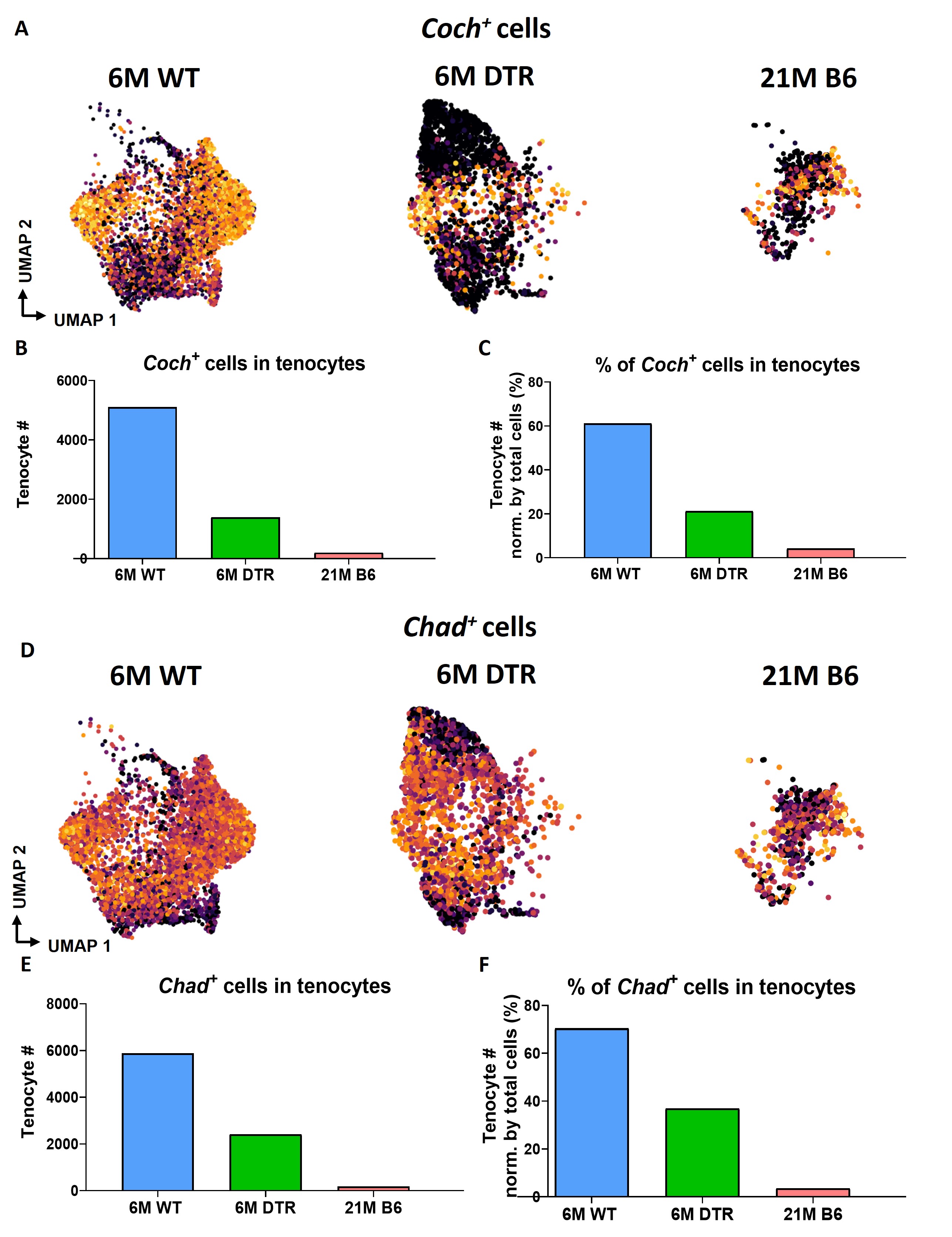

### Supplemental figure 6

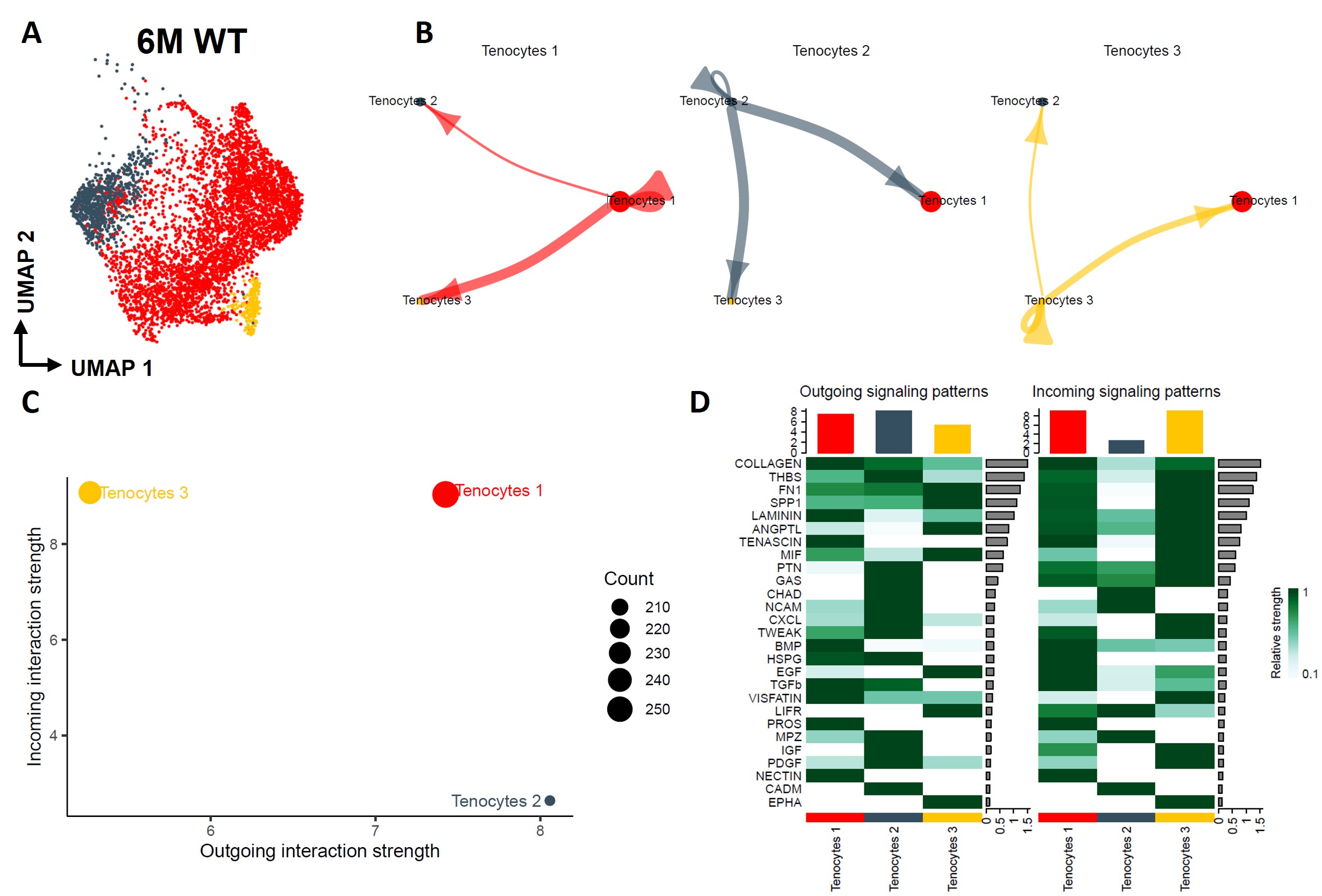

### Supplemental figure 7

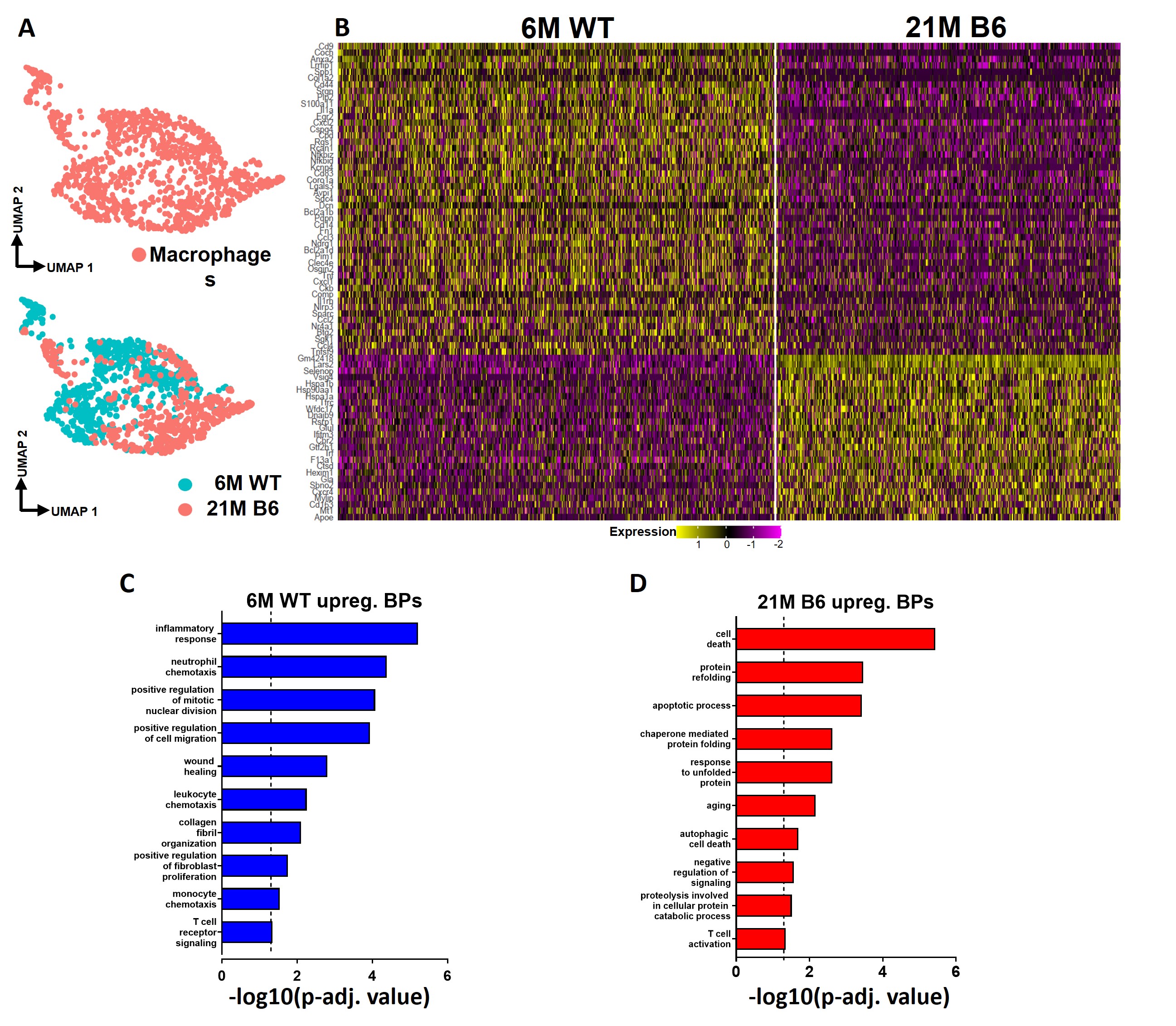
